# Inhibitory and excitatory cFos engram neurons are preferentially reactivated by sharp wave ripples

**DOI:** 10.1101/2024.12.17.628897

**Authors:** M. H. Javed, E. M. Robles-Hernandez, R. Patel, A.L. Flores Camacho, L.J. Nessler, M.G. Haberl, S. Viana da Silva

## Abstract

Sharp wave ripples (SWRs) reactivate hippocampal activity patterns during memory consolidation, but whether they recruit cFos engram cells is unclear. We tagged CA1 neurons activated during sequential exploration of two spatial contexts and later identified them *in vivo* by optotagging. We found that the cfos engram population is composed of place cells, non-place principal cells and inhibitory interneurons. cFos-tagged place cells were reused across context exposures while maintaining context specific maps. During post-behavior rest, cFos-tagged place cells and interneurons showed stronger SWR recruitment than untagged cells, with cFos identity being more closely associated with SWR modulation than firing rate features. cFos-tagged interneurons were speed- and theta-modulated, preferentially coupled to cFos-tagged place cells during SWRs, and preferentially recruited from parvalbumin and somatostatin expressing classes. After exposure to contexts outside the original tagging experience, enhanced interneuron recruitment was attenuated, whereas tagged place cells retained elevated SWR coupling. Thus, SWRs engage excitatory and inhibitory components of a cFos-tagged spatial ensemble, linking cellular engram identity to hippocampal network dynamics during memory consolidation.

## Introduction

How experiences are converted into durable memories remains a central question in neuroscience. In the hippocampus, spatial experience has been studied from two complementary perspectives. At the level of ongoing spatial representation, place cells provide spatial information and together form a map of the surrounding environment^1,2^. At the level of memory allocation, an engram can be defined as a neuronal ensemble recruited during an experience that contributes to later memory expression. In hippocampus, this causal link has been most clearly established for cFos-defined ensembles, in which cFos-expressing neurons were found to be required for memory retrieval and sufficient to trigger memory-related recall when reactivated^3–7^. Recent work has shown that cFos-tagged CA1 ensembles include place cells^8–10^, and that Fos-expressing CA1 neurons can form reliable and long-lasting spatial maps of familiar environments^11^. Moreover, cells tagged during one context can remain context-specific when animals are later re-exposed to the same context or introduced to a different environment^9^. However, the encoding of spatial experiences often unfolds across multiple related episodes rather than within a single isolated environment. Consistent with memory allocation models, distinct contextual experiences encoded close in time can recruit overlapping CA1 ensembles, thereby linking separate memories through shared neuronal populations^12^. This raises the question of whether the cFos-tagged CA1 place cell population recruited during a temporally linked sequence of spatial contexts are reused across those different context representations, and whether such reuse reflects a single stable spatial map or context-specific remapping.

At the network level, high frequency oscillations (150 - 250 Hz) in the hippocampus called sharp wave ripples (SWRs) are known to reactivate neuronal sequences of place cells during offline periods^13–17^ and are critical during the post-learning phase, as their disruption interferes with memory consolidation^18–22^. SWRs reactivate neurons that previously participated in an experience^13,23,24^ but also indicate future trajectories and emerging remapping^25,26^. Mechanistically it is thought that the repeated co-activation during SWRs promotes synaptic plasticity mechanisms required for the consolidation^27–29^. Even though SWRs are thought to play this pivotal role during consolidation, their relevance beyond a coordinated replay of trajectories of spatially tuned cells remains much less well understood. For example, it remains unclear whether particular memory encoding subpopulations of neurons, like engram cells, are recruited during post-encoding rest periods by SWRs.

Hippocampal GABAergic interneurons add a further unresolved dimension to this question. Inhibitory interneurons regulate temporal precision, movement-related firing and the synchronization of hippocampal network oscillations, including theta activity and SWRs^30–34^. SWRs are therefore not only replay events of principal-cell assemblies, but network events whose emergence and temporal organization depend on coordinated excitation and inhibition. Consistent with this, distinct CA1 interneuron classes show different speed modulation, theta phase preferences and SWR-related firing patterns^35–38^. Yet, hippocampal engram studies thus far have focused mainly on principal cells and only recent work has begun to reveal that inhibitory neurons can also form part of activity-defined memory ensembles, opening an emerging field of interneuron engram biology^39^. However, whether CA1 interneurons are recruited into cFos-defined ensembles after spatial experience, which interneuron classes are involved, and whether cFos-tagged interneurons participate in the same SWR events as tagged principal cells remain open questions.

Here, we combined cFos-dependent tagging, in vivo extracellular recordings, local field potential recordings and direct optotagging in dorsal CA1. Mice explored two distinct spatial contexts in sequence and then revisited the initial context, allowing us to tag neurons recruited during a temporally linked spatial experience across environments. We then recorded tagged and untagged cells during later re-exposure to the same contexts, post-behavior rest and rest sessions following exposure to contexts outside the original tagging experience. This design allowed us to relate cFos-defined engram identity to place-cell reuse across context representations, to the physiological and molecular identity of tagged interneurons, and to the recruitment of excitatory and inhibitory tagged cells during SWRs.

## Results

### Sequential spatial exploration induced cFos expression in principal cells and interneurons of hippocampal CA1

To examine how temporally linked spatial experiences are represented by cFos-expressing CA1 neurons, we combined cFos-tTA mice with optotagging of cFos-tagged cells using an AAV-TRE-ChR2-mKate2 viral vector (**Fig. 1a,b**). The experiment was designed to tag the collective experience of sequential exploration of two distinct spatial contexts and the revisit to the initial context, to understand if and how cFos-tagged place cells are reused across sequential spatial experiences. Mice were first habituated (around 5 days) to handling, the transfer into a different environment and the recording procedure, while doxycycline was present in the diet, suppressing TRE dependent ChR2-mKate expression. Doxycycline was then removed to open the tagging window. Approximately 48 h after DOX removal, mice explored two novel environments Context A, Context B, and were then returned to Context A (A’), after which doxycycline was restored to suppress further tagging (**Fig. 1a**). Thus, the tagged engram population corresponds to a physical neuronal trace associated with an experience, i.e. cells recruited during the sequential A-B-A’ context experience, rather than to a single location within one environment. In the following days, extracellular unit activity and local field potentials were recorded from dorsal CA1 during pre-behavior rest, re-exposure to Context A, Context B, and A’, and post-behavior rest, followed by optotagging sessions. Post hoc histology confirmed mKate expression in dorsal CA1 (**Fig. 1c**) and allowed reconstruction of tetrode bundle locations (**Fig. S1a**). We next classified recorded units as putative principal cells or putative interneurons. To do so, we first identified excitatory and inhibitory monosynaptic interactions from cross-correlograms (**Fig. 1d**) and used these labels to train a Gaussian mixture model based on the three features that best separated the ground-truth classes: firing rate, autocorrelogram mean, and spatial coverage (**Fig. 1e** and **Fig. S1b**). To identify optotagged units *in vivo*, reflecting cFos-dependent ChR2 expression during the tagging window, we delivered 473 nm light pulses at the end of each recording session. In control stimulation sessions light responsiveness was tested at 561 nm (**Fig. 1f**). Since light activation can in principle evoke both direct ChR2 mediated spiking and indirect / secondary responses, we used two steps in our optotagging analysis. First, light responsive units were identified with the Stimulus Associated spike Latency Test (SALT^40^**; Fig. S2a**). Second, light responsive units were further classified according to the latency of their first spikes (within 15 ms) after light pulse onset (**Fig. 1g-h** and **Fig. S2b**). Spike latency was defined as the mode of the first spike latency across trials, because including all spikes or using mean latency can be skewed by later secondary spikes (see **Fig S2d-e**). Light responsive units with first spike latencies within 7 ms (similar to Lakunina et al., 2026^41^) were classified as directly optotagged cells (hereafter referred to as cFos-tagged cells), whereas cells with longer latency responses were classified as putative indirect and excluded from the cFos-tagged population (**Fig. 1h** and **Fig. S2**). For compactness, figure panels use cfos+ and cfos- to denote cFos-tagged and untagged units, respectively. Across 11 optotagged mice, we recorded 1007 units in the Context A-B-A’ and associated rest sessions, of which 124 were identified as directly cFos-optotagged and the remaining 883 as not cFos-tagged. Of these recorded units, 899 were classified as putative principal cells and 108 as putative interneurons. Within the cFos-tagged population, 77.4% of cells were putative principal cells and 22.6% were putative interneurons, whereas the untagged population contained 90.9% putative principal cells and 9.1% putative interneurons (**Fig. 1i**). Thus, sequential spatial exploration recruited a cFos-tagged CA1 population composed of both principal cells (**Fig. 1j**) and interneurons (**Fig. 1k**), with inhibitory interneurons forming a substantial fraction of the cFos-tagged engram neurons. This electrophysiological classification of inhibitory cells was further supported by immunostaining for mKate and GAD67, which showed that cFos-tagged mKate+ cells included both GAD67+ inhibitory interneurons and GAD67-putative excitatory principal cells (**Fig. 1l** and **Fig. S1c**). This finding motivated us to subsequently analyze not only cFos-tagged principal cells but also cFos-tagged interneurons, which have thus far received little attention.

**Fig. 1.**
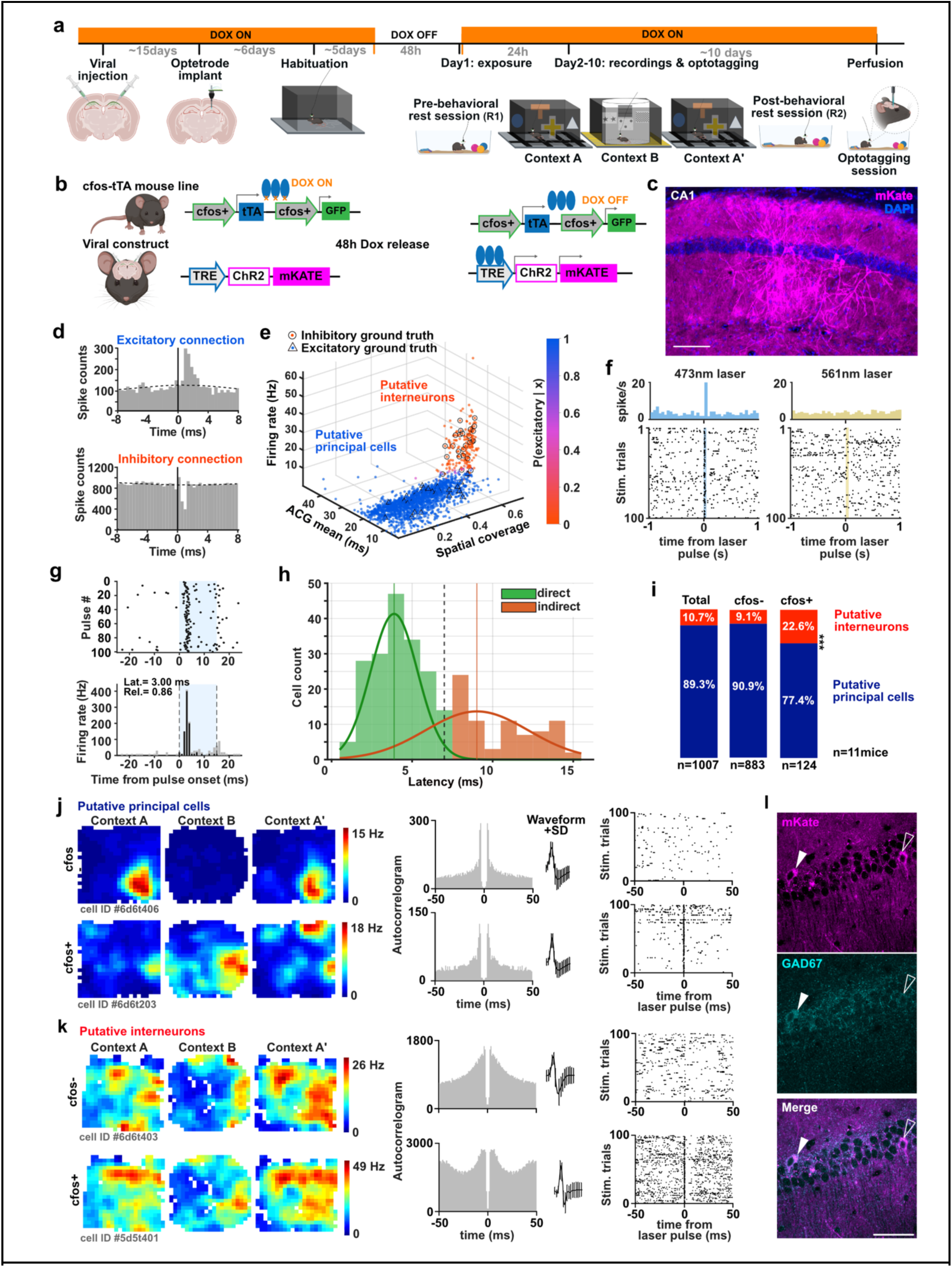
Sequential spatial exploration induces cFos expression in CA1 principal cells and interneurons. (a) Experimental timeline. cFos-tTA mice were injected with AAV-TRE-ChR2-mKate2 in dorsal CA1 and implanted with optetrodes. After habituation, doxycycline was removed from the diet to open the cfos tagging window. Mice explored Context A, Context B, and the revisit to Context A (A’) during the tagging window, after which doxycycline was restored. In the following days, mice were recorded during pre behavior rest (R1), sequential exposure to Context A, Context B, and A’, and post behavior rest (R2), followed by optotagging sessions and post hoc histology. (b) Schematic of the cfos-tTA/TRE tagging strategy. Under DOX ON conditions, cfos driven tTA is suppressed and ChR2-mKate expression is prevented. After DOX removal, cfos expressing cells activate TRE dependent ChR2-mKate expression, allowing optical identification of cFos-tagged cells *in vivo*. (c) Example image of mKate expressing cells in CA1 after spatial context exposure. (d) Examples of cross correlograms used to identify putative monosynaptic interactions and define ground truth excitatory and inhibitory units for cell type classification. (e) Gaussian mixture model classification of recorded units into putative principal cells and putative interneurons based on firing rate, autocorrelogram mean, and spatial coverage. Ground truth excitatory and inhibitory units identified from cross correlograms are overlaid. (f) Example optotagging response of a cFos-tagged unit to 473 nm laser stimulation, with no corresponding response to 561 nm control stimulation. (g) Example light-evoked responses from SALT-positive units used for latency estimation. Raster plots and peri-stimulus time histograms are aligned to light pulse onset; the stimulation window is shaded, all spikes are shown in light gray, and first spikes used to estimate response latency are shown in black. (h) Latency distribution of light responsive units used to distinguish optotagged cfos+ cells from indirect light responsive cells. Direct responses were defined by short first spike latencies after light pulse onset, whereas responses above the 7 ms latency cutoff (dashed line) were classified as putative indirect and excluded from the directly optotagged population. (i) Fraction of putative principal cells and putative interneurons among all recorded cells, cfos-cells, and directly optotagged cfos+ cells. (j) Examples of cfos- and cfos+ putative principal cells, showing spatial firing maps across Context A, Context B, and A’, autocorrelograms, spike waveforms, and optotagging responses. (k) Examples of cfos- and cfos+ putative interneurons, shown as in (j). (l) Example immunostaining for mKate and GAD67 in CA1, showing that cFos-tagged mKate+ cells include GAD67+ interneurons as well as GAD67-putative principal cells. Scale bars: (c, l) 50µm. For compactness, figure panels use cfos+ and cfos- to denote cFos-tagged and untagged units, respectively.

### cFos-tagged place cells are reused when encoding spatial representations across sequential context exposures

Out of 766 putative principal cells, 10.7% were identified as cFos-tagged and the remaining 89.3% as untagged cells. Consistent with previous work showing that cFos-expressing CA1 engrams include spatially tuned neurons^8–10^, a large fraction of cFos-tagged putative principal cells were classified as place cells. Specifically, 89.6% of cFos-tagged principal cells were place cells, compared with 75.3% of untagged principal cells (**Fig. 2a**), indicating a significant overrepresentation of place cells among cFos+ neurons relative to the overall principal cell population (binomial test, p = 0.0022). cFos-tagged place cells had higher average firing rates (p < 0.0001), higher temporal peak firing rates (p = 0.0063), and also higher burst indices (p =0.0020) than untagged place cells (**Fig. 2b**; Mann Whitney or MW test; see Supplementary Table 1 for more details on the statistics performed). cFos-tagged place cells also showed higher spatial peak firing rates (p =0.0297) than untagged place cells, while they had larger place fields (p =0.0007) with lower spatial information content (bits/spike) on average across the three Contexts (p < 0.0001). cFos-tagged place cells had a larger proportion of multi-place field cells, with two or more place fields (**Fig. 2c**; Chi-square p = 0.0101). Thus, even though cFos-tagged principal cells were more likely to code for space than untagged principal cells, the cFos-tagged place cells carried less precise spatial information, consistent with previous findings^9^.

**Figure 2.**
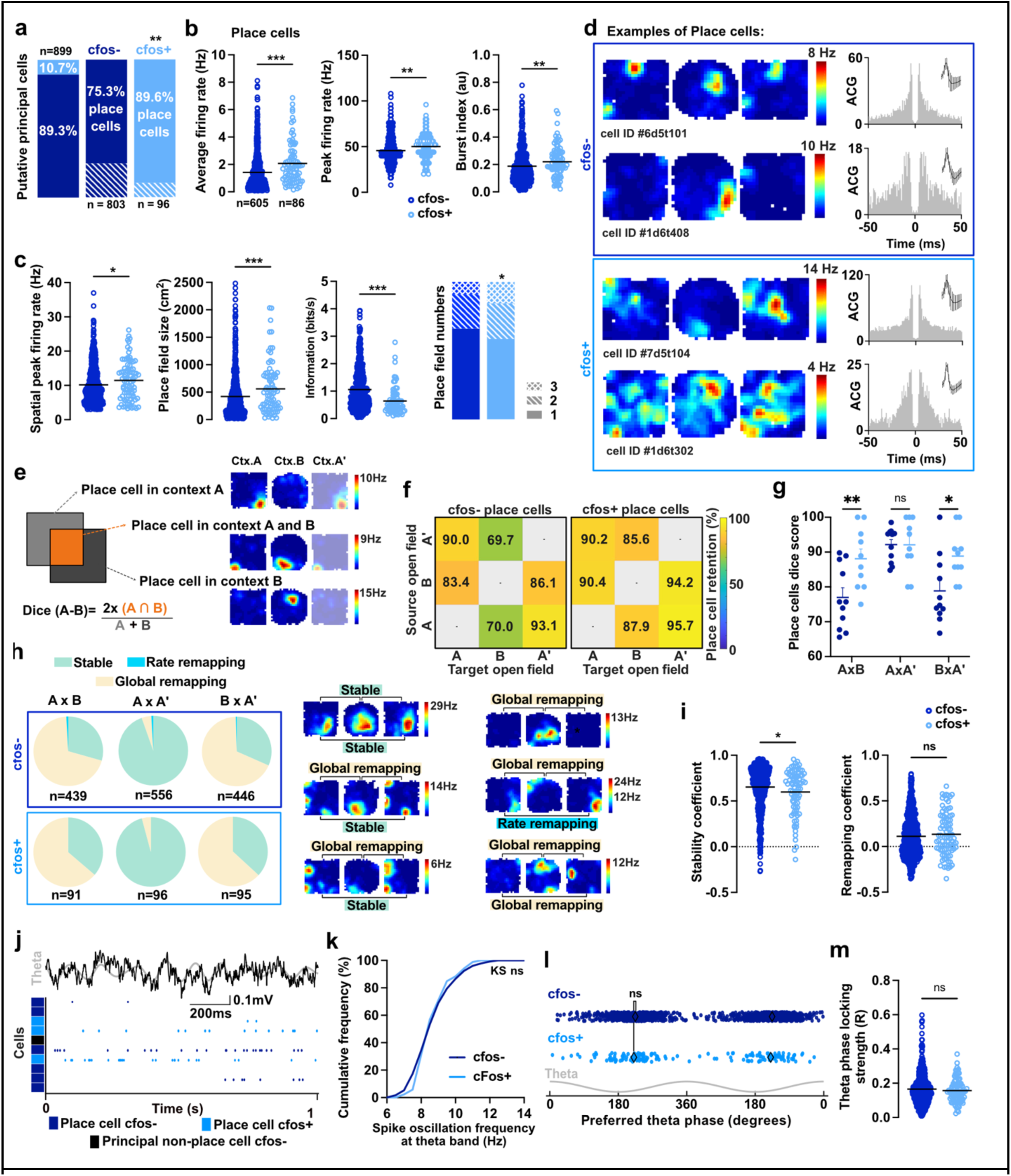
cFos-tagged place cells are reused across sequential context exposures while maintaining context-specific spatial representations. (a) Ratio of cFos-tagging within principal cells, and fraction of place cells within untagged and cFos-tagged principal-cell populations. (b) cFos-tagged place cells showed higher average firing rate, higher (temporal) peak firing rate, and higher burst index compared with untagged place cells. (c) cFos-tagged place cells also had higher spatial peak firing rates, larger place fields, lower spatial information and a larger fraction of place cells with more than 1 place field, compared to untagged place cells. (d) Examples of untagged and cFos-tagged place cells recorded sequentiallyin Context A, Context B, and the revisit to Context A (A’), with corresponding autocorrelograms and spike waveforms. (e) Schematic of the Dice coefficient used to quantify reuse of place-cell identity across context exposures. A high Dice score indicates that the same cells remain place cells across two exposures, independent of whether their place fields occur at the same spatial location. (f) Retention matrices for pairwise place cell identity for untagged and cFos-tagged place cells across Contexts A, B, and A’ show high retention of place cell identity across sequential exposures. (g) Pairwise Dice scores indicate that cFos-tagged place cells are more consistently reused between A × B and B × A’, whereas place cell reuse between A × A’ is very high and equal between untagged and tagged cell populations. (h) Spatial maps of place cells across contexts are classified as stable, rate remapping, or global remapping categories. Maps are predominantly stable between A and A’, but largely globally remap between distinct contexts. (i) Quantification of map similarity shows a significant difference in stability coefficient but no significant difference in remapping coefficient between untagged and cFos-tagged place cells. (j) Example of raw and theta-filtered LFP with simultaneous activity of untagged and cFos-tagged cells. (k) Spike oscillation frequencies within the theta band were similarly distributed between untagged and cFos-tagged place cells. (l) Preferred theta phase of untagged and cFos-tagged place cells was similarly distributed and (m) theta phase-locking strength did not differ between untagged and cFos-tagged place cells. In b, c, g, i, and m, circles indicate individual observations and horizontal lines indicate the mean. ns: P > 0.05, * P < 0.05, ** P < 0.01, *** P < 0.001.

Additional analyses of the broader putative principal-cell population, including principal non-place cells, confirmed higher firing rates in cFos-tagged principal cells and further characterized place-cell speed scores, place-field sizes, and theta-related properties of principal non-place cells (**Fig. S3**).

Since cfos expression was induced during a sequence of context exposures, we next asked whether cFos-tagged place cells were restricted to a single context or whether the same cells were reused as place cells across the three exposures. To quantify place cell identity independent of place field location, we calculated the Dice coefficient between the sets of cells classified as place cells in Context A, Context B, and the revisit to Context A (A’) (**Fig. 2d,e**). Both cFos-tagged and untagged place cells showed high pairwise retention of place cell identity across sequential exposures (**Fig. 2f**). Pairwise Dice scores were higher for cFos-tagged place cells in the A × B and B × A’ comparisons, whereas place cell reuse between A × A’ was similarly high for cFos-tagged and untagged populations (**Fig. 2g**). These results indicate that cFos-tagged cells are preferentially reused as place cells across sequential context exposures, rather than representing a population confined to a single arena. However, retention of place cell identity does not necessarily imply that cells maintain the same spatial map. We therefore classified pairwise map changes as stable, rate remapping, or global remapping (**Fig. 2h**). As expected for repeated exposure to the same context, maps were predominantly stable between Context A and A’. In contrast, place cells largely underwent global remapping between Context A and Context B, as well as between Context B and A’. The predominance of global rather than rate remapping between distinct context exposures indicates that CA1 represented Context A/A′ and Context B as distinct spatial maps^42,43^, while many of the same cells continued to be recruited as place cells across the sequence. Quantification of map similarity showed a significant difference in the stability coefficient between cFos-tagged and untagged place cells (p = 0.0317, MW test), whereas the remapping coefficient did not differ between groups (**Fig. 2i**). Together, these data show that cFos-tagged place cells are reused across sequential spatial experiences, while their spatial maps remain context specific.

Since spike patterns in the theta frequency range (6 to 14 Hz) have been shown to be important for memory encoding^44,45^, and cFos-tagged cells have been reported to display theta modulated bursts^9^, we examined place cell activity in relation to theta oscillations. Simultaneous LFP and unit recordings showed theta associated spiking of cFos-tagged and untagged place cells during exploration (**Fig. 2j**). The spike oscillation frequencies of CA1 place cells within the theta band were similarly distributed for cFos-tagged and untagged place cells (**Fig. 2k**). Preferred theta phase was also similarly distributed between groups (**Fig. 2l**), and theta phase locking strength did not differ between cFos-tagged and untagged place cells (**Fig. 2m**). Thus, the preferential reuse of cFos-tagged place cells across sequential context exposures was not explained by detectable differences in theta frequency, theta phase preference, or theta locking.

### cFos-tagged interneurons are a speed modulated and theta locked subset of CA1 interneurons

Since we identified that a fraction of the cFos-tagged cells were interneurons (**Fig. 1**, Examples in **Fig. 3a** and **Fig. S4a**), and since the contribution of interneurons to the hippocampal cfos expressing population has received little attention to date, we next characterized the activity profile of tagged interneurons during spatial exploration. Out of 108 putative interneurons, 28 cells were identified as cFos-tagged and 80 cells were untagged, corresponding to 26% cFos-tagged and 74% untagged putative interneurons (**Fig. 3a,b**).

**Fig. 3.**
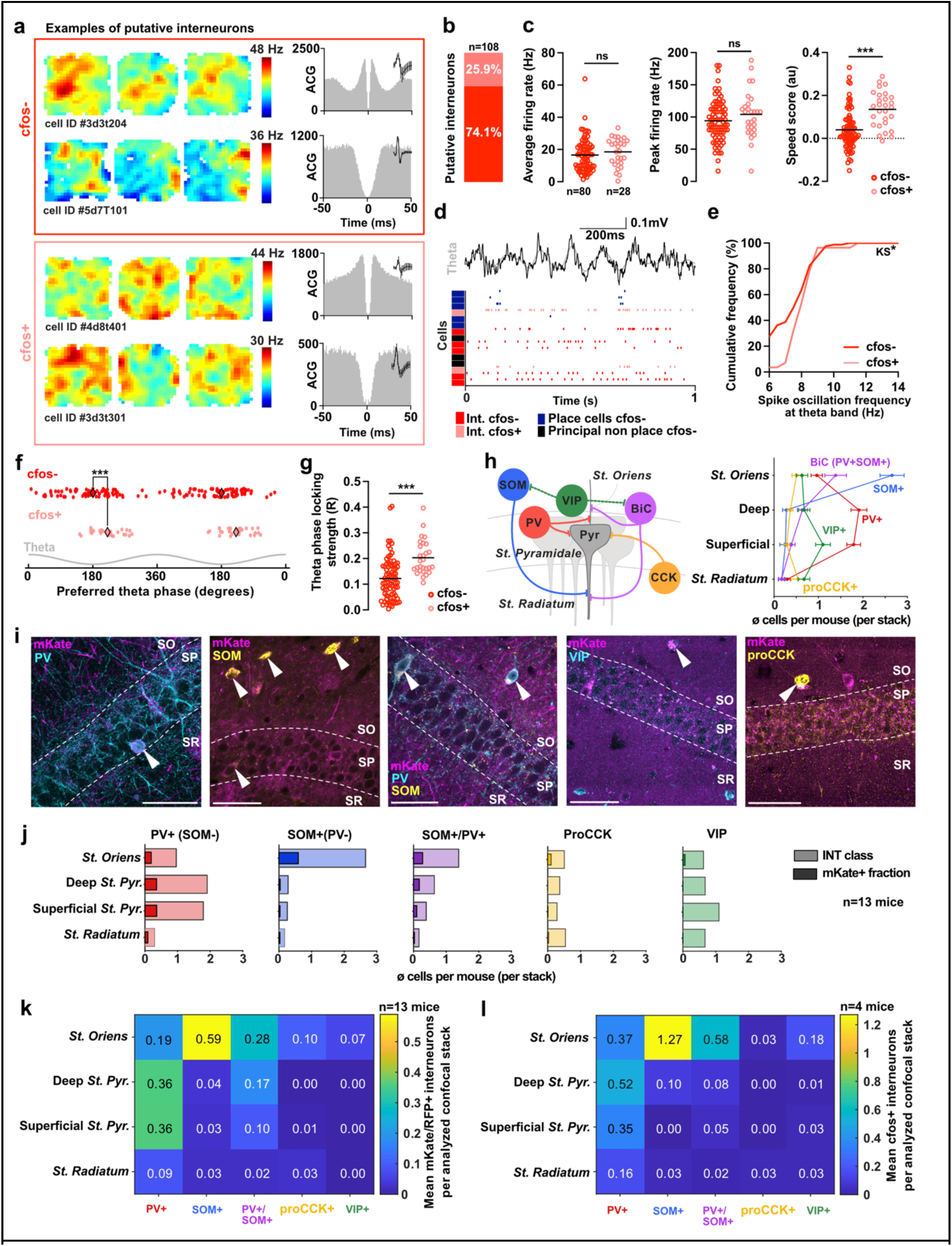
cFos-tagged interneurons are positively speed modulated, theta coupled, and include PV and SOM interneuron classes. (a) Examples of spatial firing maps across Context A, Context B, and the revisit to Context A (A’), autocorrelograms, and spike waveforms of untagged and cFos-tagged putative interneurons. (b) Fraction of cFos-tagged cells within the putative interneuron population. (c) cFos-tagged and untagged putative interneurons showed similar average firing rates and peak firing rates, whereas cFos-tagged interneurons had higher speed scores. (d) Example of raw and theta filtered LFP with simultaneous activity of untagged and cFos-tagged putative interneurons, untagged place cells, and untagged principal non-place cells. (e) Intrinsic spike oscillation frequencies within the theta band were significantly differently distributed between untagged and cFos-tagged putative interneurons. (f) Preferred theta phase of untagged and cFos-tagged putative interneurons relative to ongoing theta oscillations. (g) Theta phase locking strength was higher in cFos-tagged putative interneurons compared with untagged putative interneurons. (h) Schematic of CA1 layers and interneuron marker classes used for immunohistochemical characterization and their observed counts per layer. (i) Representative confocal images showing mKate/RFP+ cells in relation to PV, SOM, VIP and proCCK immunoreactivity across CA1 layers. Arrowheads indicate example mKate/RFP+ cells. (j) Quantification of marker-defined interneuron classes and the corresponding mKate/RFP+ fraction across CA1 layers in cFos-tTA mice. Bars show the mean number of cells per mouse per confocal stack. (k) Heat map summarizing the mean number of mKate/RFP+ interneurons per analyzed confocal stack across CA1 layers and interneuron marker classes in cFos-tTA mice. (l) Heat map summarizing the corresponding distribution of endogenous cFos+ interneurons after context exposure in naive mice. In c and g, circles indicate individual putative interneurons and horizontal lines indicate the mean. ns: P > 0.05, * P < 0.05, *** P < 0.001, Scale bars: 50μm

We first asked whether cFos-tagged interneurons differed from untagged interneurons in their general firing properties. The average firing rate and peak firing rate of cFos-tagged and untagged putative interneurons were similar (**Fig. 3c**). In contrast, cFos-tagged interneurons showed a significantly higher burst index (**Fig. S4b**; p = 0.0254, MW test) and higher speed scores (**Fig. 3c**; p < 0.0001, MW test) than untagged interneurons. Thus, cFos-tagged interneurons were not simply defined by higher overall firing rates, but were preferentially recruited among interneurons whose firing was positively coupled to the animal’s running speed during spatial exploration.

Since CA1 interneuron activity is strongly organized by theta oscillations during movement and spatial encoding, we next examined theta related firing properties of cFos-tagged and untagged putative interneurons^31^. Simultaneous LFP and unit recordings showed theta associated spiking of both, untagged and tagged interneurons during exploration (**Fig. 3d**). In contrast to the place cell population, the intrinsic oscillation frequencies within the theta band were significantly differently distributed between untagged and cFos-tagged putative interneurons (**Fig. 3e**; p = 0.0160, KS test). Moreover, the cFos-tagged population lacked units without a detectable intrinsic theta oscillation frequency, which were present among untagged interneurons. While both populations displayed organized preferred theta phase firing relative to ongoing theta oscillations, this was more coherent for cFos-tagged interneurons (**Fig. 3f**; p = 0.0038 Kuiper perm. test), which also showed stronger theta phase locking than untagged interneurons (**Fig. 3g**, p < 0.0001, MW test). Together, these findings indicate that cFos-tagged interneurons form a behaviorally and theta coupled subset of CA1 interneurons, rather than representing a random sample of the interneuron population.

Given that different CA1 interneuron classes have been shown to differ in their speed modulation, theta rhythmicity, theta phase preference, and network state dependent recruitment, the physiological differences we observed between cFos-tagged and untagged putative interneurons motivated us to determine whether cFos-tagged interneurons corresponded to specific interneuron classes. We therefore performed immunohistochemical characterization of cFos-tagged cells across CA1 layers using interneuron subtype markers for parvalbumin (PV), somatostatin (SOM), procholecystokinin (proCCK), and vasoactive intestinal peptide (VIP) (**Fig. 3h-i**). Quantification across cFos-tTA mice showed that mKate/RFP+ interneurons were detected among PV+, SOM+ and PV+/SOM+ interneuron classes, whereas overlap with proCCK+ and VIP+ cells was low or absent (**Fig. 3j,k**).

To test whether this pattern was specific to the cFos-tTA tagging strategy, we further examined endogenous cFos expression in naive animals after context exposure. These experiments showed a similar pattern, with endogenous cFos expression most evident among PV+ and SOM+ interneurons and little or no overlap with proCCK+ or VIP+ interneurons (**Fig. 3l** and **Fig. S5–S6**). Thus, context exposure recruits a selective subset of CA1 interneurons into the cFos-expressing population, including PV+ and SOM+ interneurons, while proCCK+ and VIP+ interneurons are largely excluded.

Taken together, cFos-tagged interneurons did not differ from untagged interneurons by overall firing rate, but were distinguished by positive speed modulation, a distinct intrinsic theta frequency distribution, stronger theta phase locking, and include PV+ and SOM+ interneuron classes. These findings suggest that cfos expression in CA1 interneurons reflects a selective recruitment of specific interneuron populations during spatial encoding, rather than a general consequence of high interneuron firing rate.

### cFos-tagged place cells are preferentially recruited during SWRs

SWRs provide brief time windows in which spatially distributed place cell activity can be reactivated in a compressed manner (**Fig. 4a**). We therefore next asked whether cFos-tagged place cells, which were recruited during sequential context exposure, were preferentially reactivated during SWRs in the subsequent rest period. Heat maps of the average firing rate of each unit across the SWR time windows showed that most place cells were preferentially active during the SWR time window (**Fig. 4b**). However, the probability distributions of SWR-associated population activity showed a significantly higher spiking probability of cFos-tagged place cells around SWR detection (**Fig. 4c,** p < 0.0001, KS test). To quantify if individual place cells are more recruited during SWRs, we used complementary measures of the SWR-associated firing of individual (cFos-tagged and untagged) units, across all detected SWRs (**Fig. 4d**). For individual units we computed SWR firing rate increase (SFI), the symmetric SWR modulation index (SSMI), the number of spikes per participated SWR (SpPR; **Fig. 4e**; examples shown in **Fig. S7a**). Consistently, cFos-tagged place cells showed higher SFI, higher SSMI, and more SpPR than untagged place cells (**Fig. 4f**; p < 0.0001, MW test). Moreover, the percentage of SWR participation (PSP) showed that cFos-tagged place cells were re-activated by a larger fraction of SWRs (**Fig. 4g**; p < 0.0001, MW test). Thus, cFos-tagged place cells were preferentially recruited during SWRs, both by firing more strongly during the SWRs in which they participated and by participating in more SWR events across the session. Additional analyses of non-place principal cells with SWR-aligned firing patterns and recruitment metrics showed also a preferential recruitment by SWRs for cFos-tagged non-place cells (**Fig. S7b-f**), suggesting that the higher reactivation is not dependent on the spatial coding of the cells in the environments.

**Fig. 4.**
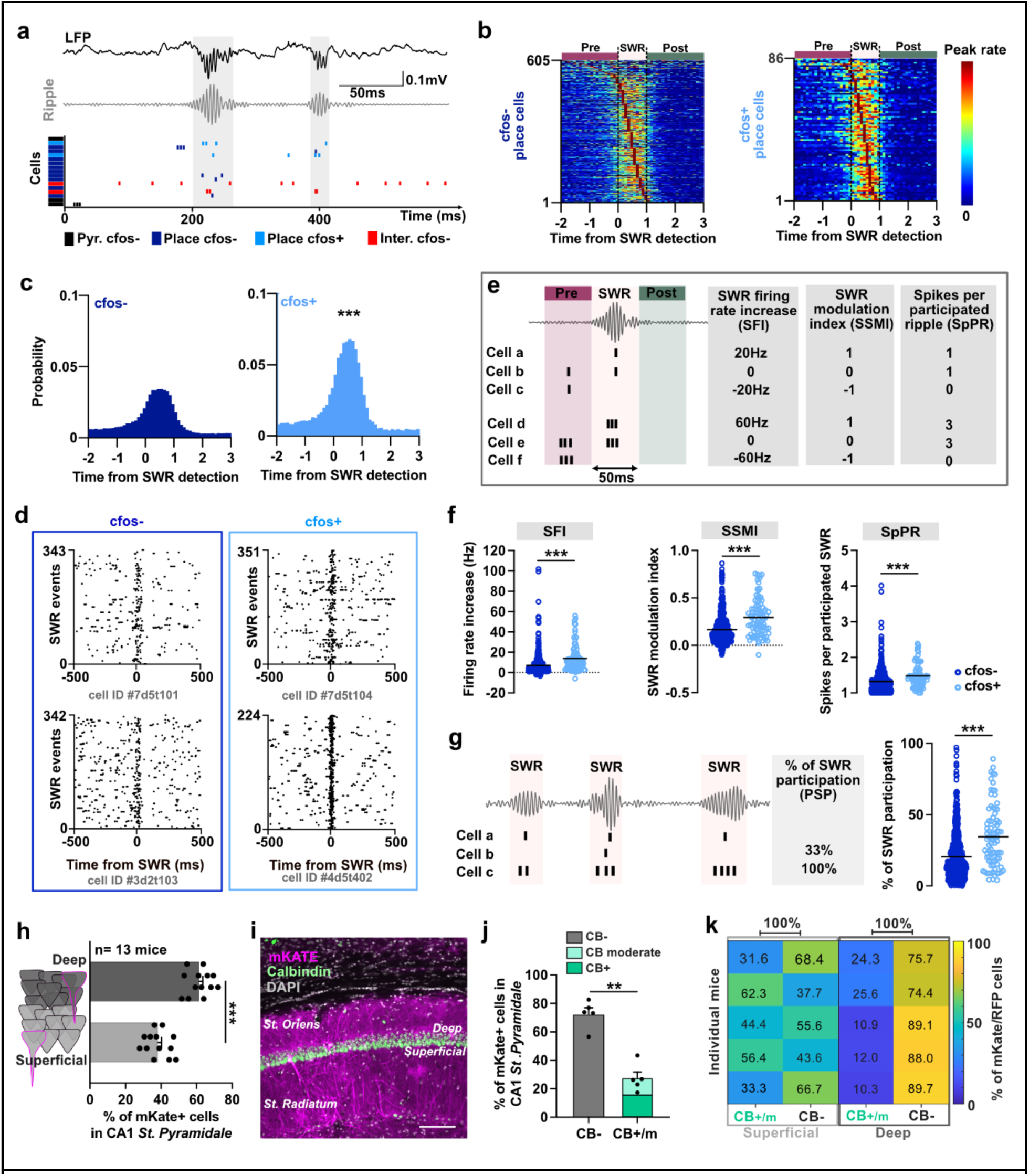
cFos-tagged place cells are preferentially recruited during sharp wave ripples. (a) Example of local field potential (LFP), ripple band filtered LFP, and simultaneous unit activity during sharp wave ripple (SWR) events. (b) SWR aligned firing rate heat maps of individual untagged and cFos-tagged place cells during post behavior rest, sorted by the relative time of their peak firing rate. Rows indicate individual place cells and columns indicate time relative to SWR detection (relative time as multiples of SWR duration). (c) Probability distributions of SWR-associated population activity shows higher spiking probability of cFos-tagged place cell firing around SWR detection compared with untagged place cells. (d) Example raster plots of individual untagged and cFos-tagged place cells across detected SWR events, centered at SWR detection. (e) Exemplary schematic illustrates different metrics used to quantify SWR recruitment of individual units. SWR firing rate increase (SFI) was calculated as the firing rate during the SWR window relative to the firing rate before the SWR window. The symmetric SWR modulation index (SSMI) quantifies relative activation or suppression during the SWR window. Spikes per participated ripple (SpPR) quantify the number of spikes during SWRs in which the cell participated. (f) cFos-tagged place cells showed higher firing rate increase (SFI), higher symmetric SWR modulation index (SSMI), and more spikes per participated SWR (SpPR) than untagged place cells. (g) Schematic and quantification of percentage of SWR participation (PSP), showing that cFos-tagged place cells also participated in a larger fraction of detected SWRs than untagged place cells. (h) Quantification of the location of mKate+ cells within St. Pyramidale of CA1 showed more cells located in deep than in superficial CA1, (i) distinguished by Calbindin (CB) staining. (j) Quantification by their CB status shows that most mKate+ cells in CA1 St. Pyr are CB-negative (moderate and strong staining of CB+ were stacked together). (k) Heatmap of individual mice shows the percentage distribution of mKate/RFP+ cells across superficial and deep CA1 CB+/intermediate and CB-groups, with the clearest preference in deep to be CB-cells. In f and g, circles indicate individual place cells and horizontal lines indicate the mean, h and j each dot represents the average per mouse. *** P < 0.001, **** P < 0.0001.

Earlier studies have described differences in the recruitment by SWRs between superficial and deep CA1 pyramidal cells, identified by Calbindin (CB) staining^46,47^. We therefore next asked whether the cFos-tagged engram cells were preferentially allocated within a specific radial or CB-defined CA1 subpopulation. Quantification of mKate+ (anti RFP stained) cell location within CA1 stratum pyramidale showed that more tagged cells were located in deep than in superficial CA1 (**Fig. 4h,I**; p < 0.0001, MW test). Classification by CB immunoreactivity further revealed that most mKate+ cells in the CA1 pyramidal layer were CB-negative, whereas moderately and strongly CB-positive cells constituted a smaller fraction (**Fig. 4j**; p = 0.0079, MW test). Consistent with the known radial organization of CB expression, strongly CB-positive mKate/RFP+ cells were enriched in superficial CA1, whereas CB-negative mKate/RFP+ cells were enriched in deep CA1 (**Fig. S7g**). Since not all superficial cells are CB-positive and not all deep cells are CB-negative we examined both the radial position and CB status across individual mice. mKate/RFP+ cells in deep CA1 were preferentially CB-negative, while superficial tagged cells showed no clear CB-positive bias (**Fig. 4k**). Complementary normalization across all radial/CB groups confirmed that tagged cells were most enriched among deep CB-negative pyramidal cells (**Fig. S7h**). Thus, the cFos-tagged CA1 engram was preferentially allocated to CB-negative cells in the deep pyramidal cell layer. This contrasts with a recent study in which repeated appetitive contextual learning preferentially recruited cFos-expressing Calbindin1-positive superficial CA1 pyramidal cells and increased their population coupling^48^. Together, these findings argue against a model in which cFos-defined CA1 ensembles are invariably allocated to a single radial pyramidal-cell subtype, and instead suggest that cFos allocation across CA1 sublayers is shaped by behavioral demand, learning history, and network state.

### cFos-tagged interneurons are strongly recruited by SWRs and coupled to cFos-tagged place cells

Since interneurons are central regulators of SWR timing and synchrony, and since cFos-tagged interneurons displayed distinct speed and theta related properties during exploration, we next examined whether they were also preferentially recruited during SWRs. SWR aligned firing rate maps showed that cFos-tagged interneuron activity was strongly concentrated around SWR detection, whereas untagged interneurons displayed more heterogeneous peri SWR firing profiles (**Fig. 5a,b**). This was also apparent when spike occurrence was quantified across pre-SWR, SWR, and post-SWR windows: whereas only 47% of untagged interneurons reached their peak firing during the SWR window, 96% of cFos-tagged interneurons peaked within the SWR window (**Fig. 5c**; p < 0.0001, Chi-Square). In agreement with this, SWR-associated population activity distributions showed a stronger enrichment of cFos-tagged interneuron firing around SWR detection (**Fig. 5d,e**; p <0.0001, KS test). Quantification confirmed that cFos-tagged interneurons had higher SWR firing increase (SFI), higher symmetric SWR modulation index (SSMI), more spikes per participated SWR (SpPR), and participated in a larger fraction of SWRs than untagged interneurons (**Fig. 5f,g**). Notably, none of the cFos-tagged interneurons showed negative SWR modulation (SSMI < 0), whereas negative modulation was observed in a subset of untagged interneurons, further indicating that the tagged interneuron population was biased toward SWR-associated activation rather than suppression. We then asked whether cFos-tagged interneurons were recruited together with cFos-tagged place cells during SWRs. SWR associated coupling analysis showed stronger coupling between cFos-tagged place cells and cFos-tagged interneurons than between cFos-tagged place cells and untagged interneurons (**Fig. 5h**; p = 0.0078, WCx signed-rank test). This indicates that cFos-tagged interneurons are preferentially co-recruited with cFos-tagged place cells during SWRs, supporting the idea that the cFos-tagged interneuron population is embedded in the same SWR associated reactivation events as the tagged spatial engram.

**Fig. 5.**
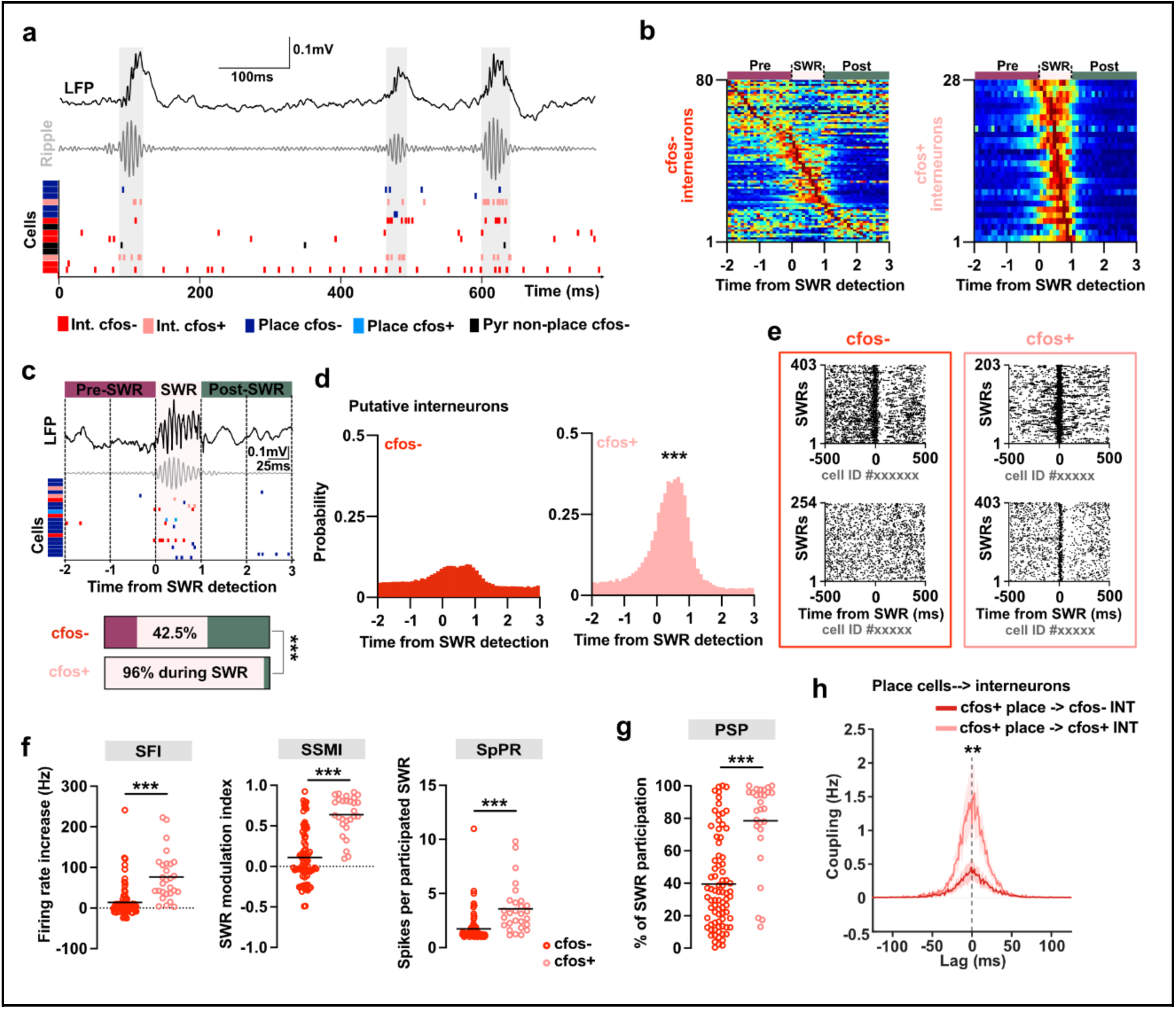
cFos-tagged interneurons are strongly recruited by SWRs and preferentially coupled with cFos-tagged place cells. (a) Example of LFP, ripple band filtered LFP, and simultaneous unit activity during SWR events. Spikes from untagged and cFos-tagged putative interneurons, untagged and cFos-tagged place cells, and untagged putative principal non place cells are shown. (b) SWR aligned firing rate heat maps of individual untagged and cFos-tagged putative interneurons during post behavior rest. Rows indicate individual interneurons and columns indicate time relative to SWR detection. (c) Example of peri SWR spiking and quantification of spike distribution across pre SWR, SWR, and post SWR windows. cFos-tagged interneuron activity was strongly concentrated within the SWR window compared with untagged interneurons. (d) Probability distributions of SWR aligned spikes show stronger concentration of cFos-tagged interneuron firing around SWR detection compared with untagged interneurons. (e) Example raster plots of individual untagged and cFos-tagged interneurons across detected SWR events, centered at SWR detection. (f) cFos-tagged interneurons showed higher SWR firing increase (SFI), higher SWR modulation index (SSMI), and more spikes per participated SWR (SpPR) than untagged interneurons. (g) cFos-tagged interneurons participated in a larger fraction of detected SWRs than untagged interneurons. (h) SWR associated coupling between cFos-tagged place cells and untagged or cFos-tagged interneurons. Coupling was stronger between cFos-tagged place cells and cFos-tagged interneurons than between cFos-tagged place cells and untagged interneurons, indicating preferential co-recruitment of cFos-tagged principal cells and interneurons during SWRs. Statistics calculated from animal-level coupling bias in the −10 to +10 ms center window. In f and g, circles indicate individual interneurons and horizontal lines indicate the mean. In h, lines indicate the mean and shaded areas indicate the error range. ** P < 0.01, *** P < 0.001.

### Post-encoding SWR participation metrics strongly distinguish cFos engram cells

To test whether SWR recruitment was associated with previously tagged cFos identity independent of their general firing properties in the environments, we used a two-block logistic-regression model. The model was fit separately for non-place principal cells, place cells and putative interneurons from the tagging room, with optotagged cFos+ identity as the binary outcome. One predictor block contained firing-rate-related features, including mean firing rate, peak firing rate, spatial firing rate and burst index. The second block contained SWR recruitment features, including percentage of SWR participation (PSP), modulation index (SSMI), SWR firing increase (SFI), spikes per participated ripple (SpPR; **Fig. 6a**; Statistical summary in Supplementary Table 4). This framework allowed us to ask whether SWR-related activity explained cFos+ identity beyond differences in overall excitability, which would be reflected in firing rate and burstiness.

**Fig. 6.**
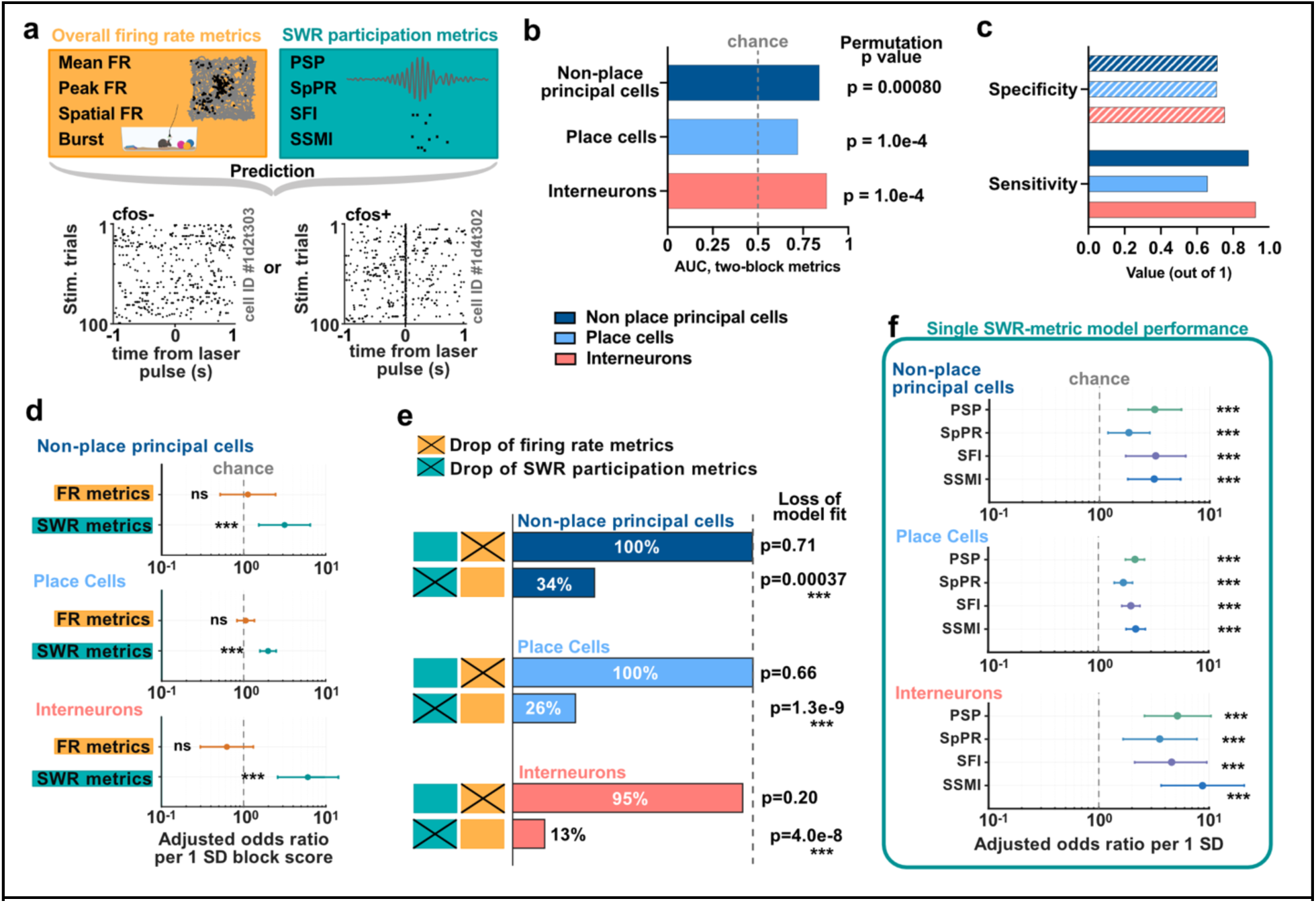
SWR participation predicts previously tagged cFos engram cell identity beyond firing rate features. (a) Schematic of the two-block logistic-regression framework used to predict true optotagging status. The firing-rate block included four metrics: mean firing rate, temporal peak firing rate, spatial peak firing rate, and burst index. The SWR participation block included four metrics: percentage of SWR participation (PSP), spikes per participated ripple (SpPR), SWR firing increase (SFI), and symmetric SWR modulation index (SSMI). (b) AUC of the full two-block model for non-place principal cells, place cells, and putative interneurons. Dashed line indicates chance-level discrimination. (c) Specificity and sensitivity at the model-selected ROC threshold. (d) Adjusted odds ratios for each predictor block per 1 SD increase in block score. Within the full model, the SWR participation block was significantly associated with true optotagging status, whereas the firing-rate block was not. (e) Predictive performance retained after removing either the firing-rate block or the SWR participation block, expressed as percentage of full-model deviance explained. Lower retained deviance indicates a larger unique contribution of the removed block. P values indicate likelihood ratio tests comparing the full model with the corresponding reduced model. Likelihood ratio tests showed that removing the firing-rate block did not significantly reduce model fit in any cell type, whereas removing the SWR participation block significantly reduced model fit in all three cell types. (f) Single-metric logistic-regression analysis of individual SWR participation metrics. For odds-ratio plots in d and f, points show odds ratios per 1 SD increase in the corresponding block score or individual metric, horizontal bars indicate 95% confidence intervals, and P values are two-sided Wald tests against OR = 1. Ns P>0.05, ** P < 0.01, *** P < 0.001.

The full two-block model distinguished cFos-positive from cFos-negative cells in all three cell classes, with high discrimination for non-place principal cells and putative interneurons and moderate discrimination for place cells (**Fig. 6b**). Discrimination was quantified by the area under the receiver-operating characteristic curve (AUC), where 0.5 indicates chance-level classification and 1 indicates perfect separation. Model AUCs were 0.846 for non-place principal cells, 0.712 for place cells and 0.865 for putative interneurons. Sensitivity and specificity at the selected model threshold showed that this performance was not driven by detection of only one class, despite the unequal numbers of cFos-positive and cFos-negative cells (**Fig. 6c**). Thus, combined firing-rate and SWR-recruitment features contained information about prior cFos tagging across both principal-cell and interneuron populations.

We next asked which feature block carried this predictive information. Block-level adjusted odds ratios showed that the SWR recruitment block was positively associated with cFos+ identity across cell classes (**Fig. 6d**). This association was significant for non-place principal cells (OR = 3.20, p < 0.001), place cells (OR = 1.97, p < 0.001) and interneurons (OR = 5.23, p < 0.0001). In contrast, the firing-rate block showed no significant association in any cell class (non-place principal cells: OR = 1.10, p = 0.818; place cells: OR = 0.99, p = 0.964; putative interneurons: OR = 0.67, p = 0.257). Thus, the statistical evidence in the two-block model was carried primarily by SWR recruitment rather than by firing-rate features.

Consistent with this interpretation, block-drop analysis showed that removing the firing-rate block retained nearly all of the full-model predictive performance, whereas removing the SWR recruitment block strongly reduced the retained deviance explained (**Fig. 6e**). This effect was especially clear for place cells and putative interneurons, where models lacking SWR recruitment metrics retained only a small fraction of the full-model performance. These results indicate that firing-rate metrics alone were insufficient to explain cFos+ identity, whereas SWR recruitment metrics accounted for most of the predictive signal.

Finally, we compared individual SWR features in single-metric logistic models to determine if individual aspects of SWR recruitment were more strongly associated with cFos-tagged identity (**Fig. 6f**). All SWR recruitment metrics showed positive associations with cFos+ identity across cell classes (interneurons and principal cells), indicating that cFos-tagged cells were distinguished by a broader SWR recruitment phenotype rather than by one uniquely informative metric. Complementary single metric AUC analyses confirmed that each individual SWR participation metrics carried discriminative information for cFos-tagged identity, whereas firing rate related metrics showed weaker or less consistent performance (**Fig. S8**). These results suggest that cFos-tagged cells are not simply high-firing cells. Rather, their identity is linked to a broader SWR recruitment profile, including how often they participate in SWRs and how strongly they fire during the SWRs in which they participate.

Together, this analysis supports the conclusion that SWR engagement is a physiological signature of previously tagged cFos cells that extends beyond firing-rate differences. This strengthens the interpretation that cFos-tagged principal cells and interneurons are preferentially embedded in SWR-associated reactivation events, rather than being identified merely because they have higher baseline excitability or greater overall firing rates.

### cFos-tagged SWR recruitment is strongest after the tagged experience and differs between place cells and interneurons

To determine whether enhanced SWR recruitment reflected a general property of cFos-tagged cells or was related to the environment in which the cells were originally tagged, we compared SWR associated activity following context exposure in the tagging room versus the control rooms that were not part of the tagging experience. Mice were recorded during rest sessions associated with the tagged Contexts A, B, and A’, and on separate days they were recorded during rest sessions associated with either a familiar control Context C or a novel control Context D (**Fig. 7a,b**). We quantified cFos-tagged versus untagged differences using Cliff’s delta across several complementary SWR metrics, including spikes per participated ripple, percentage of SWR participation, SWR firing increase, and the symmetric SWR modulation index (**Fig. 7c**). Positive Cliff’s delta values indicate stronger SWR recruitment in cFos-tagged cells.

**Fig. 7.**
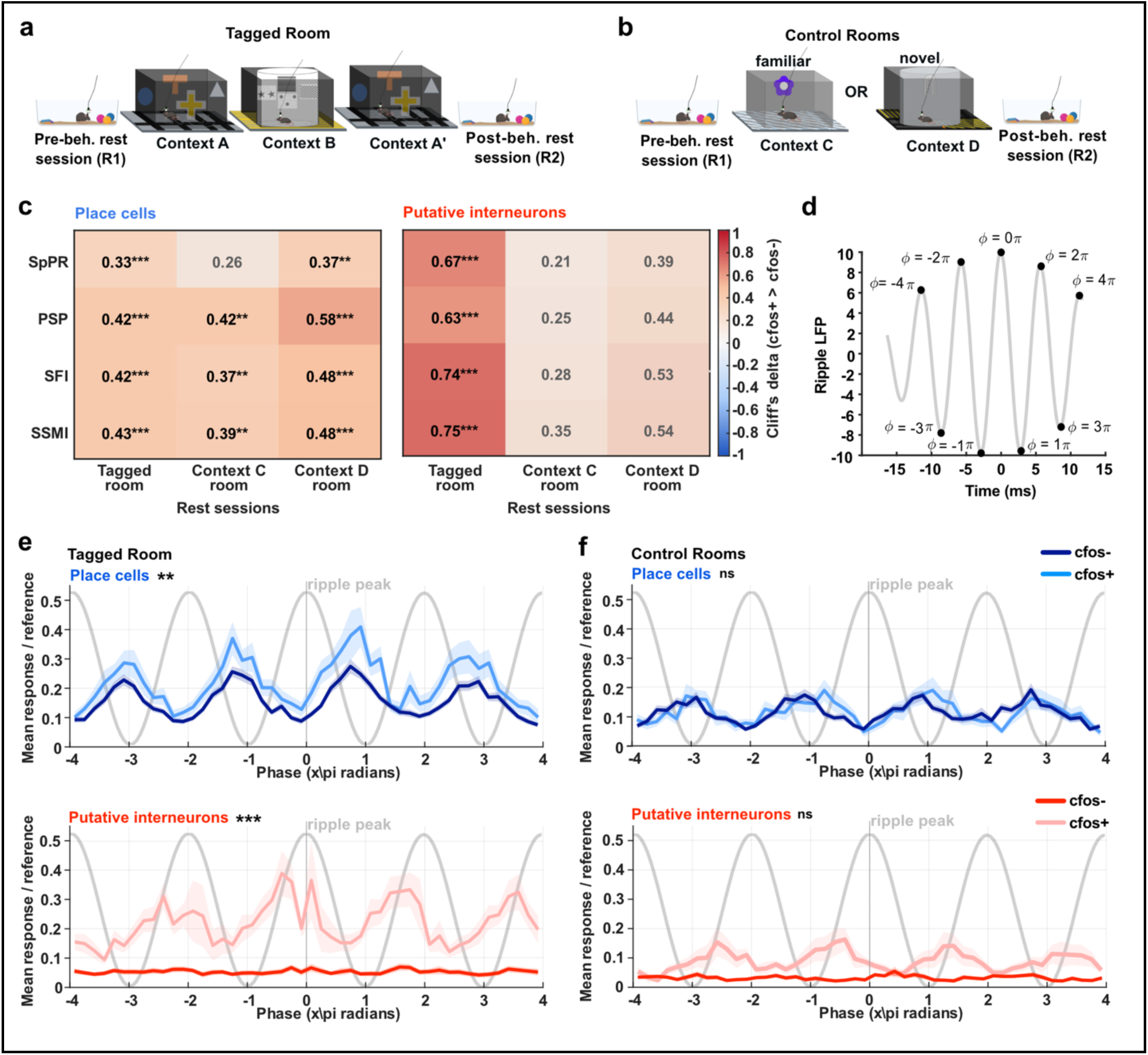
cFos-tagged place cells and interneurons show context dependent and phase organized recruitment during SWRs. (a) Schematic of recordings in the tagged room. Mice were recorded during pre behavior rest, sequential exposure to the tagged Contexts A, B, and A’, and post behavior rest. (b) Schematic of control room recordings. In separate sessions, mice were recorded during pre behavior rest, exposure to a familiar control Context C or a novel control Context D, and post behavior rest. (c) Heat maps summarizing the effect size of cFos-tagged versus untagged SWR recruitment across tagged and control room rest sessions. Values indicate Cliff’s delta, with positive values (red) corresponding to larger SWR associated values in cFos-tagged cells than in untagged cells. color intensity reflects the magnitude of Cliff’s delta, with non-significant comparisons shown at reduced opacity. cFos-tagged place cells showed positive SWR recruitment across tagged and control room sessions, whereas cFos-tagged putative interneuron recruitment was strongest and statistically significant across different metrics in the tagged room and reduced in control rooms. (d) Schematic of ripple phase assignment from the ripple band filtered LFP. Phase was expressed relative to the ripple cycle, with the ripple peak centered at 0. (e) Phase resolved SWR recruitment of untagged and cFos-tagged place cells and putative interneurons during tagged room rest sessions. cFos-tagged place cells and cFos-tagged putative interneurons showed elevated phase locked recruitment, with distinct phase profiles across the ripple cycle. (f) Same as in (e), but for rest sessions associated with control room exposure. Phase locked separation between cFos-tagged and untagged cells was reduced in control rooms, particularly for putative interneurons. Lines indicate the mean and shaded areas indicate the error range. * P < 0.05, ** P < 0.01, *** P < 0.001.

For place cells, cFos-tagged cells showed positive effect sizes across the tagged room and both control room conditions, indicating that cFos-tagged place cells retained an elevated propensity to participate in SWRs even outside the original tagging environments. However, for putative interneurons, cFos-tagged versus untagged differences were strongest in the tagged room and were reduced in the control rooms, where the effect sizes did not reach the same statistical significance. Thus, cFos-tagged place cells appear to carry a more general increase in SWR recruitment, whereas cFos-tagged interneurons show a more context-dependent recruitment profile that is most apparent during rest associated with the tagged experience.

### cFos-tagged place cells and interneurons are recruited at distinct phases of the ripple cycle

We next asked whether the elevated SWR recruitment of cFos-tagged cells was uniformly distributed across the ripple event or organized by the intrinsic phase of the ripple oscillation. To address this, we assigned spikes to the phase of the ripple band filtered LFP around SWR events (**Fig. 7d**) and compared phase resolved recruitment of cFos-tagged and untagged place cells and putative interneurons. During rest, after exploration of the context environments in the tagging room, cFos-tagged place cells and cFos-tagged interneurons both showed elevated ripple phase related recruitment compared with untagged cells, but their profiles were not identical (**Fig. 7e**). During the ascending phase of the ripple cycle, cFos-tagged interneuron recruitment increased while cFos-tagged place cell activity was reduced. Conversely, during the descending phase, cFos-tagged interneuron recruitment decreased while cFos-tagged place cell recruitment increased. This complementary phase organization suggests that cFos-tagged interneurons are not merely more active during SWRs, but are recruited at specific phases of the ripple cycle where they may help structure or counterbalance the timing of cFos-tagged place cell activity.

After exploration of the context in the control rooms, phase-related separation between cFos-tagged and untagged cells was markedly reduced, particularly for putative interneurons (**Fig. 7f, Fig. S9**). This indicates that the strong phase organized recruitment of cFos-tagged interneurons is preferentially expressed during SWRs associated with the tagged experience. Together, these results suggest that cFos-tagged place cells retain elevated SWR coupling across contexts, whereas cFos-tagged interneurons are preferentially recruited in a context specific and ripple phase dependent manner. This supports the idea that tagged interneurons participate in the organization of SWR reactivation related to the tagged spatial experience, rather than simply reflecting a globally more excitable interneuron population.

## Discussion

In this study, we examined the functional properties and participation in network events of cFos expressing hippocampal engram cells, which were tagged during context encoding. Sequential spatial exploration recruited a CA1 population composed of both principal cells and inhibitory interneurons. Within the principal cell population, cFos-tagged cells were frequently place cells, which were reused across the sequential context exposure which happened within a short time frame. However, reuse of place cell identity did not imply reuse of the same spatial map: cFos-tagged and untagged place cells remapped between distinct contexts, while re-exposure to the same context was more stable. Thus, cFos-tagged cells repeatedly participated in spatial representations, while their spatial tuning remained context specific.

This distinction between cellular reuse and map stability is important for interpreting the tagged population. The A-B-A’ protocol was designed to tag a temporally linked spatial experience rather than a single environment. The high Dice scores indicate that many of the same cells remained place cells across exposures, whereas the remapping analyses show that CA1 still represented the contexts as distinct spatial maps. Together with their larger place fields and lower spatial information, cFos-tagged place cells may therefore provide a reusable cellular substrate for spatial experience, while the precise place field pattern remains flexible. This indicates that the tagged engram is not a single fixed map of one arena.

A central finding is that cFos-tagged cells were preferentially recruited during SWRs. This was evident for tagged place cells and for tagged interneurons, and was captured by several SWR metrics including modulation, firing increase, spikes per participated ripple, and percentage of ripple participation. Importantly, SWR recruitment was not simply explained by firing rate features. A two block model showed that SWR participation metrics accounted for tagged identity beyond firing rate related measures. This supports the idea that SWR recruitment is a physiological signature of cFos-tagged cells, rather than a trivial consequence of higher baseline firing or general excitability.

These results provide a direct link between two levels of memory research that have often been studied separately: molecular or cellular engram tagging and network level SWR reactivation. Our data does not show that SWRs are required for cFos allocation or memory consolidation, because this study did not perturb SWRs or tagged cells. It does, however, show that previously tagged cells are preferentially embedded in SWR events during later rest. This provides a plausible mechanism by which cells that were active at different locations or different moments of an extended experience could be recruited within compressed network events. Such SWR related co recruitment may help link distributed components of a spatial experience into a coherent ensemble.

The identification of cFos-tagged interneurons extends this link beyond principal cells. Tagged interneurons were not simply the highest firing interneurons: their average and peak firing rates were similar to those of untagged interneurons, whereas they were distinguished by positive speed modulation, a distinct intrinsic theta frequency distribution, stronger theta phase locking, and strong SWR recruitment. Immunohistochemical analyses further indicated that the tagged interneuron population includes PV and SOM interneurons, whereas proCCK and VIP interneurons were rarely tagged. Similar endogenous cFos patterns in naive animals after the same context exposure argue that this is not merely an artifact of the cfos-tTA optotagging strategy or mouse line, but reflects the intrinsic cfos expression after the encoding of novel environments. Thus, context exploration recruits a selective subset of interneurons into the cFos-expressing population.

The SWR coupling between tagged interneurons and tagged place cells suggests that these inhibitory cells are not only cFos-tagged, but that they are embedded in the reactivation dynamics. Tagged interneurons were more strongly recruited during SWRs, participated in a larger fraction of SWR events, and showed stronger SWR associated coupling with tagged place cells than did untagged interneurons. Their phase relation to tagged place cells was also distinct: tagged interneuron recruitment increased during phases in which tagged place cell activity was relatively reduced, and decreased when tagged place cell recruitment increased. This complementary phase organization suggests that tagged interneurons may help balance network excitability and structure the timing of tagged principal-cell activity during SWRs. This interpretation is consistent with evidence that CA1 PV+ interneurons can organize post-learning hippocampal network oscillations and stabilize neuronal ensembles required for memory consolidation^49^, while our data further suggest that a cFos-tagged subset of interneurons is selectively embedded in SWR-associated reactivation after spatial context exposure.

However, our control context data refine this interpretation. Here cFos-tagged place cells retained elevated SWR coupling even after exposure to control environments, suggesting that cFos-tagged principal cells either have a broader propensity to participate in SWRs or that they distinguish to a lesser extent between the tagging and the control environment. In contrast, tagged interneuron recruitment was strongest after re-exposure to the tagging environments and reduced in control contexts. This indicates that the interneuron component of the tagged engram is more context dependent than the place cell component. Rather than just reflecting a globally more excitable or SWR coupled interneuron class, the tagged interneuron recruitment appears to be preferentially expressed during SWR events associated with the original tagged experience.

We expect that future experiments combining cell type specific tagging, time-resolved tagging in different environments, targeted perturbation, and context specific replay decoding will help to resolve now emergent questions of the excitatory and inhibitory engram balance and whether and how tagged interneurons are causally involved in organizing SWR reactivation and memory expression, and if individual interneuron types are assigned to different engrams.

Together, our findings show that cFos-tagged CA1 ensembles are not restricted to principal cells, but include selective inhibitory interneuron populations. Both excitatory and inhibitory tagged cells are preferentially recruited during SWRs, with interneurons showing context dependent and phase organized coupling to tagged place cells. These results support a model in which SWRs provide a network state that links distributed components of a cFos-tagged spatial experience, while inhibitory engram cells help shape the timing and context specificity of this reactivation.

## Methods

### Mice

All experiments were approved by the Berlin Landesamt für Gesundheit und Soziales (LAGeSo), and followed the German animal welfare act and the European Council Directive 2010/63/EU on protection of animals used for experimental and other scientific purposes. All optotagging experiments were performed in cFos-tTA mice (RRID:IMSR_JAX:018306) that were purchased and bred at the Charité animal facility. For these *in vivo* recordings 13 mice were used across 2 cohorts, 3 of these were male (∼12 months old, ∼35 g) and 10 were female (∼12 months old, ∼28 g). Two animals out of 13 were excluded from the analysis of the *in vivo* data (leaving a final number of 11 mice), since the viral injection did not match the tetrode location and therefore no opto-tagged cells were detected in these two mice. These mice were still included for immunohistochemistry quantifications since the bilateral viral injections were successful, and they underwent the exact same protocol as the other mice. All mice were provided with water and regular food, ad libitum. Three to four weeks before the start of the experimental paradigm mice were switched to doxycycline-containing diet (40mg/Kg, (DOX-ON, Ssniff Spezialdiäten GmbH, Germany). Upon surgery, mice were individually housed within a temperature and humidity-controlled chamber Zoonlab Uniprotect NG M (U5-MFR) with a 12-hour light/dark cycle.

Additionally, 4 male C57BL/6J mice were used for histological control experiments to assess endogenous cFos expression in different interneuron classes after context encoding. These mice did not undergo viral injections nor optetrode implantation. They were otherwise handled equally to the cFos-tTA mice, habituated to the experimenter, room and transfer into other environments. They were sacrificed 2 hours after the exposure to the novel environments, which consisted of the Context A, Context B, Context A’ with the pre- and post-rest sessions.

### Viral vector injection

Mice were injected bilaterally in the dorsal CA1 with AAV-TRE-ChR2-mKate2, an adeno-associated viral vector (AAV), expressing channelrhodopsin2 (ChR2) and the mKate2 fluorescent protein under the control of the tetracycline-responsive element (TRE). The viral vector was produced by the Charité Viral vector Core Facility (VCF) and stored in a freezer at −80°C until usage. For the stereotaxic injections, the mice were first anesthetized with Isoflurane (4%) in an anesthesia chamber before being moved to the stereotaxic frame, where they were kept under constant isoflurane (1-2%) for the duration of the surgery. A glass micropipette (Blaubrand intraMARK 1 - 5 μL) was pulled using a micropipette puller (Sutter Instrument Model P-1000 Flaming) and was loaded using a 1:5 dilution of the AAV-TRE-ChR2-mKate2 viral vector diluted in sterile saline solution. 0.5 μL were infused bilaterally into the CA1 of the hippocampus (AP 1.85 mm, ML +/- 1.75 mm, DV −1.3 mm from Bregma). The mixture was infused at a rate of 70 nL / minute and was allowed to diffuse at the injection site for 10 minutes before retrieval of the pipette.

### Microdrive assembly and implant

Custom microdrives were built based on the Neuralynx Electrode Interface Boards (EIB-18) which housed 4 tetrodes (Tungsten, 0.0127 mm diameter, gold plated to 150 - 250 kΩ, California Fine Wire Company, or Platinum, 0.017 mm diameter platinum plated to 150 - 250 kΩ, California Fine Wire Company) and an optic fiber Thorlabs FP200URT (200 ± 5 μm core diameter). The electrodes were cut to a distance of <1mm from the extremity of the optic fiber and glued separately to it to form the recording bundle. The microdrive implant was implanted into the right hemisphere above the dorsal CA1 region (AP 1.85 mm, ML 1.75 mm, DV −1.3 mm from Bregma). In the subsequent days the tetrodes were lowered slowly until the pyramidal layer was reached as determined by the shape of the local field potential (LFP), specifically the presence of sharp wave ripples during rest and clear theta oscillations during mobility, and the appearance of distinct clusters of units.

### Behavior protocol, Doxycycline and ChR2 expression

cFos-tTA (cFos tetracycline-transactivator) mice were kept on a doxycycline (DOX-ON) diet for about 4 weeks to suppress expression of ChR2. During this time the mice were habituated to the room, to the handling and to the recording procedure over several days while the doxycycline was already present for 3-4 weeks in their diet. At the time the tagging window was opened, mice had therefore already been extensively habituated to handling, transfer into and out of the arena, and placement in both the arena and home cage. This ensured that the novelty experienced during the tagging window reflected the first exposure to the novel environments (Context A and Context B), rather than the procedure itself or being placed into or transferred across arenas. When tetrodes reached the CA1 pyramidal layer (identified by SWRs and Theta waves), the diet was switched to DOX-OFF (normal diet) for 48 hours thus opening the window of cfos-dependent expression^9^ of ChR2 and mKate2 and the mice remained in their familiar home cage environment during these 48 hours to avoid novelty exposure.

Then, on the day of the first exposure to the new environments, the mice were first recorded for ∼10 minutes in their home cage while resting to ensure stable readings and recordings of SWRs. Mice were exposed to Context A (square box with black walls, 50 x 50 cm with distinct floor and wall cues), followed by Context B (circle with white wall, 50 cm diameter with distinct floor and wall cues), and returned back to Context A (Context A’). The exposure to every environment lasted approximately 10-15 minutes to ensure that the mice explored each environment thoroughly to induce place fields. After exposure to these two novel environments (Context A and B), the mice were placed back in their home cage and allowed to rest for ∼20 minutes for SWRs to appear. Right after this exposure to the novel environments, the DOX-ON diet was re-introduced in the home cage to end the tagging window of cFos-expressing cells.

From the subsequent day onwards we conducted *in vivo* electrophysiology using optetrode recordings and were able to identify cFos-tagged cells expressing ChR2 (Fig. 1) in the dorsal CA1 of these mice (Fig. S1a), while they were re-exposed to the same environments (Fig. 1). At the end of Day 2 recordings through to the last day of recordings, tetrodes were lowered daily to find additional cFos cells that had been tagged on Day 1. This approach enabled longitudinal recording and identification of the original cFos expressing neurons, as the mice subsequently returned to the same two environments daily. Apart from the experimental setting, the mice were exposed only to their individual home cage.

For our first cohort (7 mice with optotagged cells), to ensure we retained the ability to optogenetically reliably identify cFos-tagged cells with light, after Day 4 of recordings, the mice were again switched to DOX-OFF for 48 hours while being maintained in their home cage, re-exposed to Context A and Context B, and then put back on the DOX-ON diet. Since the environments were the same during both tagging windows, the cells would potentially re-express cFos, or new cFos cells would encode the now familiar environments in Phase 2, or the original Phase 1 cFos cells would still carry ChR2 from Phase 1 but be tagged in Phase 2. In either case, cFos cells would represent the environments of interest and increasing familiarity. The second cohort (4 mice with optotagged cells) was not exposed to the second DOX window as it was determined that just one iteration of DOX ON/OFF was enough to express ChR2 for the duration of our experiments. Additionally, they followed an interleaved experimental design incorporating control-room exposures. In this cohort, Days 1 and 2 followed the same tagging and recording procedures as the first cohort without the additional DOX-OFF window. From Day 3 onwards, experimental and control days were strictly alternated throughout for up to two weeks. Each control day consisted of two sessions separated by 1.5 hours: a familiar control environment (Context C) and a novel control environment (Context D).

Context C was considered familiar since mice had been previously habituated for a week to the room and to an open field area with no cues during threading and recording preparations, making it a well-known setting. During experimental recordings, the arena was modified, consisting of a grey square open field with a specific floor and stable cue on the wall. In contrast, Context D was entirely novel, as the mice had never been exposed to this room or arena before; it was introduced for the first time on Day 3. For this context, the arena consisted of a circular grey open field with a distinct cue on the wall and floor, designed to be specific only to this context. For each session, mice were first placed in their home cage, then exposed to an open-field arena containing condition-specific visual and tactile cues, then returned to their home cage, and finally undergoing an optotagging session. Tagged-room and control-room sessions were recorded using identical electrophysiological and optotagging procedures.

In addition to the cFos-tTA mice, four C57BL/6J mice were used as control animals for histological comparisons. These mice did not undergo viral injections or optetrode implantation, however, they were handled and habituated to the tagging/tagged room using the same habituation procedures as the experimental mice prior to their first exposure. Environmental exposure procedures were the same as experimental mice, however they were not subjected to electrophysiological recordings or optotagging. Mice were perfused for subsequent histological analysis ∼90 minutes after the post-behavioral rest session.

### Data acquisition

Data was acquired using a DigitalLynx 4SX Recording System (Neuralynx, Bozeman, MT, USA). Mice were connected to the recording system through a custom-made multichannel head-stage preamplifier connected to multi-wire tether (Neuralynx). A series of elastics allowed to counterbalance the weight of these devices to ensure the animal could run freely. Signals were amplified (5,000-20,000 times) and band-pass filtered (0.6-6 kHz) for spike detection, and spike waveforms above a threshold of 35-50 μV were time-stamped and digitized at 32 kHz for 1 ms. LFP was recorded continuously in the 0.1-900 Hz band from one of the wires from each tetrode at a sampling rate of 32 kHz. The position of the mouse was tracked using red and green colored light emitting diodes (LEDs) with a video camera mounted above the experimental area, with an acquiring rate of 30 Hz.

### Spike Sorting

Spikes were sorted manually using a customized version of the spike sorting software MClust, MATLAB 2009b (Redish, A.D. Mclust. https://redishlab.umn.edu/mclust). Different combinations of peak amplitude, energy, and peak differences were used to identify clusters. For a given cluster, the same boundaries were used across the entire recording session, but the clustering did not include spikes that occurred during optotagging sessions to avoid bias. Clusters were excluded from analysis if they were not separable from either the noise or other single units. To ensure stability all units included in the study were required to have at least 5 spikes in both the pre- and post-behavioral resting sessions (R1 and R2), while they were not required to have spikes during the behavior blocks.

### Optotagging

Optotagging sessions were conducted every day (except after Day 1) after the second 20 minute sleep/rest session (R2). A total of 100 pulses of a 473 nm laser at an intensity of ∼10-20 mW was applied for 15 ms pulses to excite and tag cells that had expressed cFos and ChR2. The 100 pulses were set to random inter-pulse intervals between 5 and 20 seconds long to avoid synaptic potentiation. Pulse Pal v2 (Sanworks, CA, USA) was used to set all laser parameters and a MLL-FN-473-100mW frequency-stabilized slim laser (Photontec-Berlin, Germany) was used to generate light pulses.

A control laser at 561 nm (MLL-FN-561-100mW, Photontec-Berlin, Germany) was used to determine if the optotagged cells were activated by heat or light of an unintended wavelength. Each control session was set to the same parameters as the 473 nm laser and run once per mouse.

### Identification of optotagged cells

Single units were classified using SALT (stimulus-associated spike latency method), a first-spike latency test used for optogenetic identification of genetically tagged neurons^40^. For each unit, we extracted the first spike occurring within a 15 ms window after each light pulse and binned first-spike latencies at 1 ms resolution. To construct the null distribution, we sampled windows of identical duration from the inter-pulse baseline period while excluding epochs within 5 ms of laser onset and offset. The observed SALT statistic was the median Jensen-Shannon divergence between the stimulus-evoked first-spike latency distribution and bootstrapped baseline latency distributions, and significance was assessed against baseline-versus-baseline comparisons. Units with SALT-based P < 0.01 and a light-evoked spike probability of at least 0.10 spikes per TTL were classified as light responsive. To further limit the chance of including secondary (indirectly) activated units we calculated the response latency, where spikes were quantized in 1 ms steps and the latency was determined as the modal light-evoked response of the first spike within the light pulse. Due to an apparent bi-modal distribution (for visualization, the modal latency distribution was fitted with two Gaussian components; **Fig. 1h**) of the modal light-evoked response of the units, we then applied a conservative fixed latency criterion where a modal latency <= 7 ms was classified as directly optotagged, whereas responsive cells with modal latency > 7 ms were classified as putatively indirectly activated and excluded from the directly optotagged pool.

### Cell type classifications of interneurons, principal cell and place cells

A set of ground truth neurons were identified as excitatory or inhibitory neurons using the cross-correlogram activity to identify monosynaptic connections and characterize if their effect is excitatory or inhibitory towards the post-synaptic cell, using the cross-correlograms within the CellExplorer framework^50^. Spike trains were concatenated across behavioral epochs within session, and cross-correlograms were computed at 0.2-ms resolution over a 120ms time window. Candidate excitatory interactions were defined by significant causal peaks within 1.0-2.8 ms after the presynaptic spike, whereas candidate inhibitory interactions were defined by significant causal troughs within 1.0-4.0 ms. Significance was evaluated against a convolution-based predictor with Poisson confidence bounds (alpha = 0.001, convolution window 10 ms), following the convolution-based framework implemented in CellExplorer and related CCG-based approaches^50–52^. To reduce contamination by spike-overlap and shared-input artifacts, the anti-causal reference window was defined as −4 to −1 ms, and for units detected on the same Tetrode the central −1 to +1 ms interval was excluded from baseline estimation. In addition, candidate excitatory interactions in which the first significant causal deflection was an inhibitory trough followed by a later positive rebound were reassigned as putative inhibitory candidates. All candidate monosynaptic interactions were subsequently reviewed manually by experienced investigators using detector-matched correlogram visualizations and only unambiguous cases were retained, whereas any ambiguous, low-confidence interactions were discarded.

Based on the ground truth connections a Gaussian Mixture Model (GMM) was trained using the three best parameters out of twelve candidate parameters (firing rate, spike half width, peak latency, AB ratio, PV ratio, rise time, auto-correlogram (ACG) mean value, ACG tau decay, ACG tau rise, spike duration, burst index, spatial coverage), which were evaluated for their 1D GMM classification accuracy on the ground truth cell types. Features were z-scored using labeled cells only. We then fit class-conditional Gaussian models separately to excitatory and inhibitory ground-truth populations (one multivariate Gaussian per class) and computed posterior class probabilities using empirical class priors. Each neuron was assigned to the class with the higher posterior probability. The final model used the following three features: spatial coverage, firing rate, ACG mean value to classify the single units into putative principal cells or putative interneurons.

Putative principal cells were further classified as place cells if they had at least one defined place field within one of the environments recorded. Spatial maps were calculated with a bin size of 2.5cm per bin and place cells had at least a spatial firing rate above 3 Hz, and a minimum place field size of 8 bins, where the cell fired at least 30% of the cell’s peak firing rate. Remaining principal cells, which did not fulfill these criteria, were grouped as non-place principal cells.

### Mean firing rates, peak firing rate, and spatial peak firing rate

Mean firing rates during open field exploration were defined as the number of spikes per sec while the mouse was moving above a velocity of 2cm/s, to avoid periods of slow wave activity. Peak firing rates were defined as the maximum firing rate, based on 250 ms time bins whereas the spatial peak firing rate was calculated as the maximum firing rate across the spatial bins of an environment.

### Burst index and Speed score

We calculated a burst index per cell, as the ratio of the number of spikes occurring in bursts (with inter-spike interval less than 6 ms) divided by the total number of spikes. We further calculated the bursts per minute as the number of bursts during the three exposures to environments (Context A, B, A’) divided by the recording duration of the three sessions. We calculated the speed score as the Pearson’s correlation between the instantaneous firing rate and the instantaneous velocity. The instantaneous firing rate was calculated for firing rates in 250 ms time bins while the mouse was running at a velocity greater than 2 cm/s.

### Spatial coverage

Spatial coverage was used to quantify how broadly each cell’s firing was distributed across the open field. For each cell, the two-dimensional firing rate map was restricted to visited spatial bins, excluding unvisited bins marked as NaN. The remaining bin firing rates were sorted in descending order, and the cumulative sum of firing rate was normalized to the total activity across the map. Spatial coverage was then defined as the fraction of visited bins required to account for 75% of the cell’s total map activity.

Thus, low spatial coverage values indicate that most firing was concentrated in a small portion of the environment, consistent with spatially restricted firing fields. Higher values indicate that firing was distributed across a larger fraction of the environment, reflecting broader or more spatially diffuse activity. This measure is a 2D implementation, adapted from previous coverage-based descriptions of hippocampal spatial firing, including the spatial coverage and field-size measures used by Royer et al. 2010^53^, and extended here to quantify the concentration of activity in open-field rate maps.

### Spatial information (rate)

The spatial information rate and the spatial information used were calculated as described previously^54^ using probability density and firing rates at particular places in the environment relative to the overall environment. Spatial information rate in bits per second is divided by the overall mean firing rate of the cell to give the spatial information in bits/spike. It provides a measurement of how specific the spatial encoding of the cell is. A high spatial information implies a focused and concentrated firing of the cell in a particular place rather than widespread firing in larger areas.

### Dice score of place cell identity

For identified place cells, the Dice score was calculated as 2 × |A ∩ B| / (|A| + |B|), where A and B are the sets of cells classified as place cells in each context. A high Dice score indicates that the same cells remain place cells across two environments and are coding for space in both environments. This metric is intentionally designed to be independent of whether their place fields are remapping or occur at the same spatial location.

### Remapping and stability

Remapping was calculated as the correlation coefficients (Pearson’s correlation) of the rate maps between Context A and Context B, whereas stability was calculated as the correlation coefficients of the rate maps between the first visit of Context A and the revisit of Context A (Context A’). The correlation coefficients have a range between −1 and +1, where a coefficient of 0 means no correlation of activity between the two environments, whereas a positive and high coefficient (0.8 to 1) implies a strong correlation and a negative coefficient implies an inverse correlation between the rate maps.

### Global remapping versus rate remapping

For each cell, open-field rate maps were generated separately for OF1, OF2, and OF3. For each pairwise comparison (OF1-OF2, OF2-OF3, OF1-OF3), we analyzed only cells that met the place-cell criteria in both sessions. We calculated the observed Pearson correlation between the two spatial rate maps and compared it to a null distribution obtained by circularly shifting the spike timestamps within one recording and recomputing the surrogate rate map 1,000 times. Cells whose observed map correlation exceeded the shuffle-based significance threshold (p < 0.05) were classified as stable, whereas cells that did not pass this threshold were classified as global remapping. For stable cells, rate remapping was quantified as the normalized absolute change in peak firing rate, abs(rate1-rate2)/(rate1+rate2), and for globally remapping cells, field displacement was quantified as the Euclidean distance between the rate-weighted centroids of the primary place field in the two sessions.

### Theta phase locking and intrinsic theta rhythmicity

Theta phase locking was quantified during open-field recordings by assigning each spike to the instantaneous phase of the local theta rhythm. LFPs from the cell-layer channel were band-pass filtered in the theta range (6–14 Hz), z-scored, and Hilbert-transformed to extract instantaneous phase. Only spikes occurring during locomotion, defined as speed >2 cm/s, were included. For each cell, spike phases were pooled across open-field sessions. The preferred theta phase was calculated as the circular mean of the spike-associated theta phases, and phase-locking strength was quantified as the mean resultant vector length. Rayleigh’s test was used to assess non-uniformity of the phase distribution.

Intrinsic theta-band spike oscillation frequency was estimated from each cell’s spike-time autocorrelogram. Autocorrelograms were computed with 5 ms bins over a 600 ms lag window and smoothed. The power spectrum of the autocorrelogram was then calculated using a multitaper spectral estimate, and the intrinsic theta frequency was defined as the frequency with maximal power within the theta range of 6.2–14 Hz.

### SWR events

SWR events were detected using a custom-made algorithm used previously^55^. In short, LFP recordings were band-passed from 150-500 Hz, and the root mean square (RMS) was calculated using a five point (2.5 msec) sliding window. High-frequency events (putative SWR) were detected during periods of immobility (< 2cm/s) as events when the RMS signal deviated from the mean RMS by greater than 3.5 standard deviations (SD) above the baseline and have at least 4 high powered cycles. Events that occurred less than 6 msec apart were merged.

### SWR participation metrics and Symmetric SWR modulation index (SSMI)

The *SWR firing increase (SFI)* was calculated as the firing rate during SWRs minus the firing rate during the matched pre-SWR reference windows. For each cell, spikes were summed across all detected SWR intervals and divided by the total SWR duration; reference-window spikes were summed across all matched reference windows and divided by the same total SWR duration. SFI therefore captures the absolute increase or decrease in firing rate during SWRs, in Hz.

*Percentage of SWR participation (PSP)* quantified how often a cell participated in SWR events. It was calculated as the percentage of detected SWRs in which the cell fired at least one spike. *Spikes per participated ripple (SpPR)* measured the mean number of spikes emitted by a cell during a SWR in which that cell was active. For each cell, only SWRs containing at least one spike from that cell were included, and the number of spikes in those participating SWRs was averaged.

For each unit, the *symmetric SWR modulation index (SSMI)* was calculated during rest sessions as a measure of SWR-associated activity relative to a matched pre-SWR reference window. For each SWR, the reference window had the same duration as the SWR and was placed one SW duration before the detected SWR onset, leaving a gap of one SWR duration between the reference window and the SWR to minimize the risk of including spikes around the detected SWR onset. If this reference window overlapped with another detected SWR, it was iteratively shifted earlier in 25 ms steps until it no longer overlapped with any detected SWR. For each unit and SWR, a modulation index was calculated as the difference between spike counts in the SWR and reference windows, normalized by their sum plus a small numerical constant: SSMI = (nSWR - nref) / (nSWR + nref + ε), where ε = 1 × 10^-7. This constant was included only to avoid division by zero; events with no spikes in either window therefore had an SSMI of 0. The final SSMI for each unit was computed as the mean SSMI across all analyzed SWRs, so that each SWR contributed equally to the cell-level estimate. This event-wise symmetric formulation differs from a pooled SMI, in which spike counts are first summed across all SWRs and reference windows before calculating a single modulation value per cell^37^. The event-wise approach prevents rare SWR participation from producing a maximal cell-level modulation score, which can occur when a sparsely firing cell fires during only one or a few SWRs and not in the corresponding reference windows. And more generally the event-wise calculation prevents overestimating the modulation when the spike counts are low, as is often the case for place cells (**Fig. S7a**). In addition, the symmetric denominator provides a conservative event-wise estimate when cells fire in both windows, as is often the case for interneurons. In these cases, normalizing by the larger spike count alone can overstate modest SWR-related differences, whereas normalization by the total activity across both windows preserves modulation direction while reducing this bias. Positive SSMI values indicate higher firing during SWRs than during the reference window, values around 0 indicate similar activity in the two windows, and negative values indicate reduced firing during SWRs.

### Ripple Phase Analysis

To quantify the timing of spikes within the ripple cycle, LFP traces were analyzed during detected SWR events. For each SWR, the LFP was ripple-band filtered using the same filter as for SWR detection, z-scored, and transformed with the Hilbert transform to obtain the instantaneous ripple phase. Spikes occurring between the detected SWR onset and offset were assigned the phase of the nearest LFP sample. To compare phase locking across SWRs, troughs were detected in the filtered ripple waveform, and spike phases were aligned to the circular mean trough phase of each SWR. For each cell, all SWR-associated spike phases were pooled across ripples, and the mean phase angle, resultant vector length, and Rayleigh test for non-uniformity were calculated. For the cumulative phase-response plots, spike phases were also represented relative to the center ripple peak, allowing responses to be visualized across ripple cycles around the SWR center. To quantify recruitment around SWR-associated phase peaks, the center-aligned phase response of each cell was normalized by its SWR versus pre-SWR recruitment and integrated within predefined phase windows around the expected ripple-cycle peaks, yielding per-cell AUC measures that were summarized across animals and compared between groups. cFos-positive and cFos-negative cells were then compared separately by cell class and recording context.

### Interneuron coupling

SWR-associated place-cell to interneuron coupling was quantified using SWR-restricted cross-correlograms (**Fig. 5h**). cFos-tagged place cells were used as source cells, and coupling was compared between cFos-tagged interneuron targets and untagged interneuron targets in the tagged room. For each session, spike trains were restricted to detected SWR windows, and directed cross-correlograms were computed with 1 ms bins over a 250 ms window centered on the source-cell spike. Pair-level CCGs were computed as raw target-spike counts at each lag relative to source-cell spikes and were normalized by the total analyzed SWR duration. The resulting values represent an SWR-restricted pairwise co-firing rate, reported as counts per second. Pair-level CCGs were averaged within animal, and group means with SEM were plotted across animals. For statistical comparison, coupling was averaged in the center window around zero lag, from −10 to +10 ms. For each animal, we calculated the paired coupling bias: (cFos-tagged place cell -> cFos-tagged interneuron) - (cFos-tagged place cell -> untagged interneuron). The animal-level bias values were tested against zero using a one-sided Wilcoxon signed-rank test, testing the directional hypothesis if cFos-tagged place cells would show stronger coupling to cFos-tagged interneurons.

### Multi-Environment Comparisons Heatmaps

Cell-level metrics were grouped by recording context, cell type and cFos/opto-tagging status. For each metric and context, cFos-positive and cFos-negative cells were compared using a two-sided Mann-Whitney U test. Ranks were averaged in the case of ties, and p-values were computed using the normal approximation with tie correction and continuity correction. The heatmap reports the effect size as Cliff’s delta rather than the raw group mean difference. Cliff’s delta quantifies the degree of separation between two distributions. It is defined as: delta = P(cFos+ value > cFos-value) - P(cFos+ value < cFos-value). Thus, values range from −1 to +1. A positive value indicates that the metric tends to be higher in cFos-positive cells, whereas a negative value indicates that the metric tends to be higher in cFos-negative cells. A value near zero indicates similarity between the two groups. In the heatmaps, each cell shows Cliff’s delta, with significance indicated as * p < 0.05, ** p < 0.01, and *** p < 0.001. For SWR-associated metrics, R1 and R2 values were averaged at the single-cell level before group comparisons. SWR heatmaps were organized by rest context: Tagging room Rest, Context C Rest, and Context D Rest.

### Predictive value analysis of post-encoding cell metrics

To assess whether post-encoding cell properties were informative about cFos engram-cell identity, we fitted logistic regression models with cFos+/true-optotagged status as the binary outcome. Analyses were performed separately for non-place principal cells, place cells, and interneurons. Predictors were organized into two balanced metric blocks: SWR participation metrics, consisting of SSMI, SFI (Firing increase), spikes per participated ripple, and PSP (% participation); and the firing rate metrics in the second block, consisting of mean firing rate, temporal peak firing rate, spatial peak firing rate, and burst index.

For the block-level analysis, each individual metric was z-scored within the analyzed cohort. The four z-scored metrics within each block were averaged to generate one SWR participation score and one firing rate score per cell. These two block scores were then entered together into a logistic regression model. Adjusted odds ratios were reported per 1 SD increase in each block score. To quantify the relative predictive contribution of each block, we compared the full two-block model with reduced models in which either the SWR participation block or the firing rate block was removed. Loss of model fit was quantified by the increase in model deviance, likelihood-ratio testing, and the percentage of full-model deviance-explained performance retained after block removal.

Overall model performance was evaluated from the fitted probabilities using receiver-operating characteristic analysis and the area under the curve (AUC). For threshold-based classification summaries, the probability threshold was selected to maximize sensitivity minus the false-positive rate, and sensitivity, specificity, positive predictive value, and negative predictive value were reported at this threshold.

In a subsequent analysis, we assessed which SWR participation metric carried the strongest signal within each cell class. For this, SSMI, SFI, spikes per participated ripple, and PSP were each fitted as single-metric logistic regression models using common complete cases across the four SWR metrics. Single-metric AUCs and odds ratios were compared to rank the SWR participation metrics, while interpreting this comparison cautiously because the SWR-derived metrics are partially correlated (e.g. the SSMI is correlated to a composite of participation and spikes per participating SWR).

### Histology and Immunostaining

In addition to electrophysiology recordings, we performed histology and immunostainings to identify cFos expressing cells and certain interneuron types, namely Parvalbumin (PV), Somatostatin (SOM), Vasoactive intestinal peptide (VIP), and Cholecystokinin (CCK) as well as for Calbindin D28K (CB) as a marker of the superficial stratum pyramidale in CA1 or Glutamate decarboxylase 67 kDa isoform (GAD67) as a marker for GABAergic cells.

All 13 implanted cFos-tTA mice were immunostained for mKate (anti-RFP), to enhance the fluorescence signal, together with PV and SOM, or proCCK and VIP to identify major interneuron classes. Five implanted cFos-tTA mice (from the second cohort) were additionally stained for mKate together with CB and GAD67 respectively. The four C57BL/6J control mice were co-stained for cFos together with following interneuron markers: GAD67, PV and SOM, proCCK and VIP, CCK8 and SNCG.

For the cFos-tTA mice after the last session of recording and for the C57BL/6J mice 90min after the novel context exposure, the mice were injected with a solution of ketamine and xylazine that was 10% of the mouse’s body weight in mL (3 mL for a mouse weighing 30 g). The mice were transcardially perfused with 1x phosphate-buffered saline (PBS) solution followed by 4% solution of paraformaldehyde (PFA) in 1x PBS. The extracted brains were left overnight at 4°C in 4 % PFA after which they were washed in cold PBS solution. Following that, the brains were sectioned into 50 μm coronal slices using a Leica VT1000S vibratome and placed in cold PBS solution. For immunostaining, the slices were first washed 3 times in PBS for 10 minutes each. Blocking and permeabilization was performed next by incubating the slices in a mixture of PBS, 0.5 % Triton X-100, 3 % bovine serum albumin (BSA) and 4% normal goat serum (NGS) for 1 hour at room temperature (24°C). Slices were then washed 3 times in a solution of PBS and 0.5 % Triton X-100 for 10 minutes each. Slices were then incubated in a primary antibody solution containing 0.8 % Triton X-100, 3 % BSA, 8 % NGS, and PBS agitating for 48 hours at 4°C. In cFos-tTA mice, to enhance the mKate signal, sections were stained using Rabbit pan-RFP antibody (ChromoTek). Sections were stained with primary antibody markers for interneuron types, Rat anti-Somatostatin-14 (ThermoFisher MA5-16987) and Chicken anti-Parvalbumin (Synaptic Systems) or Guinea Pig anti-Calbindin D28K (Synaptic Systems 214 005) and Rabbit anti-GAD-67 (ThermoFisher PA5-21397) or Rabbit anti-proCCK (Frontier Institute Co.,Ltd. MSFR105030) and Guinea Pig anti-VIP (Synaptic Systems 443 005).

In non-implanted mice, all sections were stained with Chicken anti-cFos (Synaptic Systems 226 009), and with primary antibody markers for interneuron types such as Mouse anti-Parvalbumin (Sigma-Aldrich P3088) and Rat anti-Somatostatin-14 (ThermoFisher MA5-16987) or Rabbit anti-SNCG (Assaygenie CAB2524) and Guinea Pig anti-CCK8 (Synaptic Systems 438 004), or Rabbit anti-proCCK (Frontier Institute Co.,Ltd. MSFR105030) and Guinea Pig anti-VIP (Synaptic Systems 443 005) or Mouse anti-Beta-IV Spectrin (Sigma-Aldrich MABN1727) and Rabbit anti-GAD-67 (ThermoFisher PA5-21397) or Guinea Pig anti-Calbindin D28K (Synaptic Systems 214 005) and Rabbit anti-GAD-67 (ThermoFisher PA5-21397).

Slices were washed 3 times in a solution of PBS and 0.1% Triton X-100 for 10 minutes each. Then the slices were incubated in secondary antibody solution containing 1x PBS, 4 % NGS, and 3 % BSA with species-specific Alexa Fluor-conjugated secondary antibodies (Thermo Fisher Scientific), including goat anti-chicken Alexa Fluor 488, goat anti-rabbit Alexa Fluor 555, goat anti-rat Fluor 647, goat anti-mouse Alexa Fluor 488/647, and goat anti-guinea pig Alexa Fluor 555/647, for 24 hours at 4°C. After secondary incubation, the slices were washed 3 times 15 min in 1x PBS and stained with DAPI diluted at 1:10.000 in 1x PBS for 20 minutes and mounted using ProLong Diamond Antifade Mounting Medium.

Electrode tracks and recording locations were confirmed using PV-stained overview images, in which layers were clearly visible (**Fig. S1a**).

### Cell Quantification of immunohistology

Image stacks of immunohistochemically stained slices were acquired using a Nikon SoRa Spinning Disk Confocal CSU-W1. Confocal z-stacks (60X magnification, 0.8μm optical sections) were processed and analyzed using Fiji (ImageJ^56^) for manual cell quantification. For cFos-tTA mice, both hemispheres were analyzed, whereas for C57BL/6J mice only one hemisphere was analyzed. All quantifications were restricted to the CA1 region of the hippocampus, which was subdivided into stratum oriens (SO), stratum pyramidale (deep and superficial layers), and stratum radiatum (SR).

Neuronal activity co-localization was assessed using condition-specific activity markers: mKate in cFos-tTA mice and labeling of endogenous cFos expression in the C57BL/6J control mice. Co-localization within CA1 was manually determined by identifying cells exhibiting overlapping signals between the activity marker (cFos or mKate) and each interneuron marker (PV, SOM, proCCK, VIP) or CB.

Interneuron subtypes within CA1 were identified using their respective immunohistochemical markers and quantified independently. PV and SOM positive interneurons were categorized into three independent populations: PV single-positive (PV+), SOM single-positive (SOM+) populations, and PV/SOM double-positive (PV+/SOM+) cells, each analyzed separately across all quantifications.

The proportion of co-labeled cells was calculated as the number of interneurons co-expressing the activity marker divided by the total number of interneurons of each respective class, and expressed as a percentage. Spatial co-labeled interneuron distribution was analyzed within CA1 subregions. For this purpose, activity-labeled interneurons were quantified either relative to the total number of interneurons of each class in CA1 or normalized to the total number of co-labeled interneurons across CA1 subregions/layers. In addition, activity-normalized co-localization was calculated as the number of co-labeled interneurons divided by the total number of mKate-positive or cFos-positive cells.

Cell counts were aggregated within animals before calculating percentages, so that each animal contributed one value per analysis. This count-weighted approach estimates the distribution of the total quantified mKate/RFP-positive population within each animal and avoids giving equal weight to sections with markedly different cell yields. Animals, rather than individual sections, were treated as the experimental unit for group-level summaries.

## Acknowledgements

We thank D. Mitrovic-Tartanoglu, S. J. Swer and P. Ayyildiz for experimental assistance, and all other members of the Haberl and Viana da Silva laboratories for feedback throughout the duration of the project.

## Authors contributions

MHJ, MGH and SVS designed the study. MHJ, ERH, RP, ALFC, LJN, MGH and SVS performed experiments, MHJ, ERH, MGH and SVS analyzed data, performed statistical analysis and made figures. MHJ, ERH, MGH and SVS wrote the manuscript with comments from all the authors.

## Data and materials availability

The data are available upon request to the corresponding authors. Code is available from the corresponding authors on request and will be deposited in the lab’s Guithub repository: https://github.com/memorycircuits

