## Supplementary Figures and Material for "Inhibitory and excitatory cFos engram neurons are preferentially reactivated by sharp wave ripples"

**This PDF file includes:**

Figures. S1 to S9 (pages 2-12)

Supplementary Tables 1 to 5 for statistical analysis (pages 13 to 16)

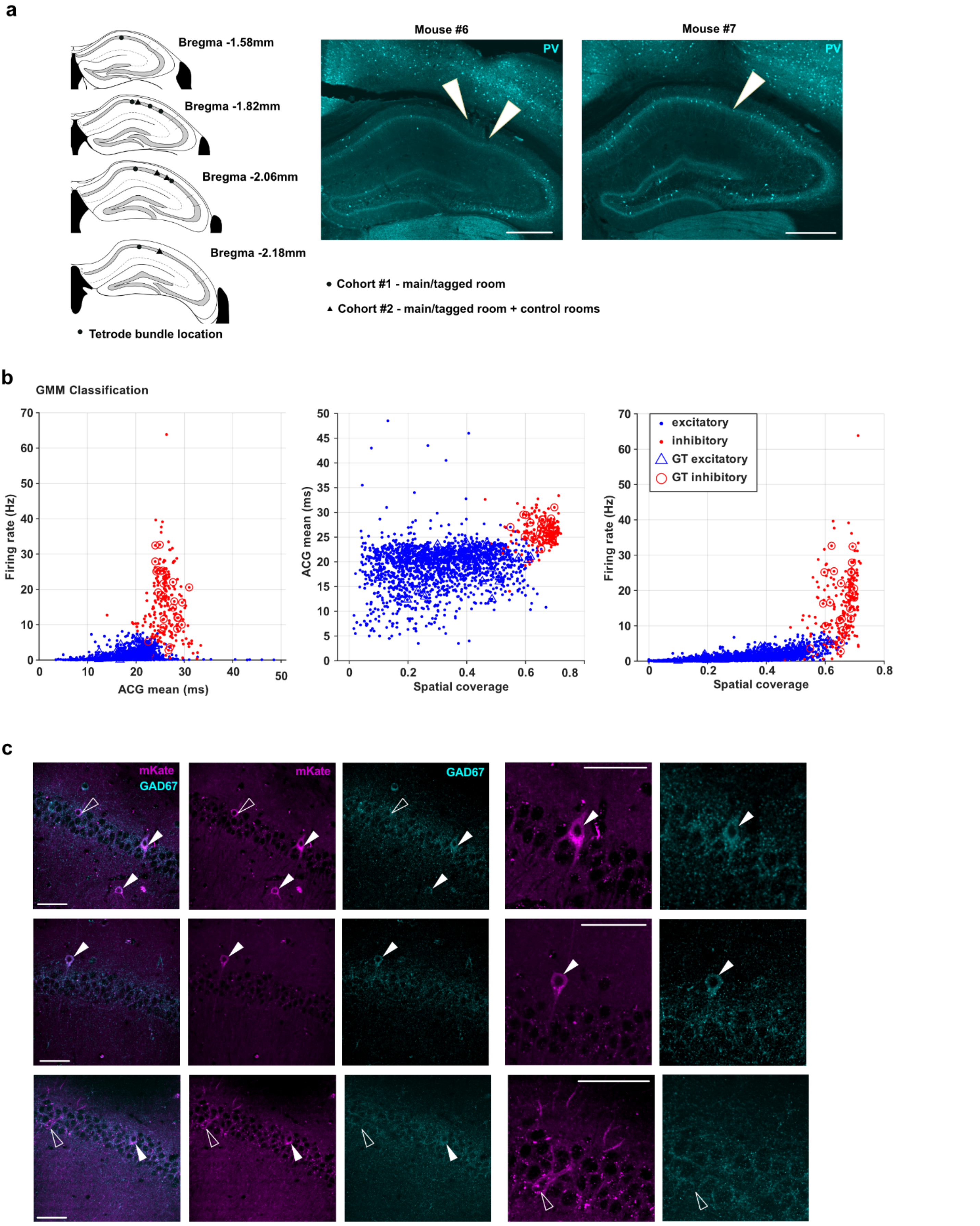


**Fig. S1. Recording locations and validation of principal cell and interneuron classification.**
(a) Anatomical locations of tetrode bundle tracks across mice, shown on coronal hippocampal sections at the indicated bregma levels. Circles indicate mice from cohort 1 recorded in the tagged environments, and triangles indicate mice from cohort 2 recorded in both tagged and control environments. Representative parvalbumin stained sections from two mice show tetrode tracks in dorsal CA1. Arrowheads indicate recording tracks. (b) Gaussian mixture model classification of recorded units into

putative principal cells and putative interneurons. Two dimensional projections show firing rate, autocorrelogram mean, and spatial coverage, with classified putative principal cells in blue and putative interneurons in red. Ground truth excitatory and inhibitory units identified from cross correlogram based monosynaptic interactions are overlaid. (c) Example confocal images show mKate and GAD67 immunostaining in CA1, supporting the electrophysiological classification by identifying mKate+ cells with and without GAD67 expression across CA1 layers. Filled and open arrowheads mark example mKate+ cells with different GAD67 labeling. Scale bars (a): 500μm (c) 50μm

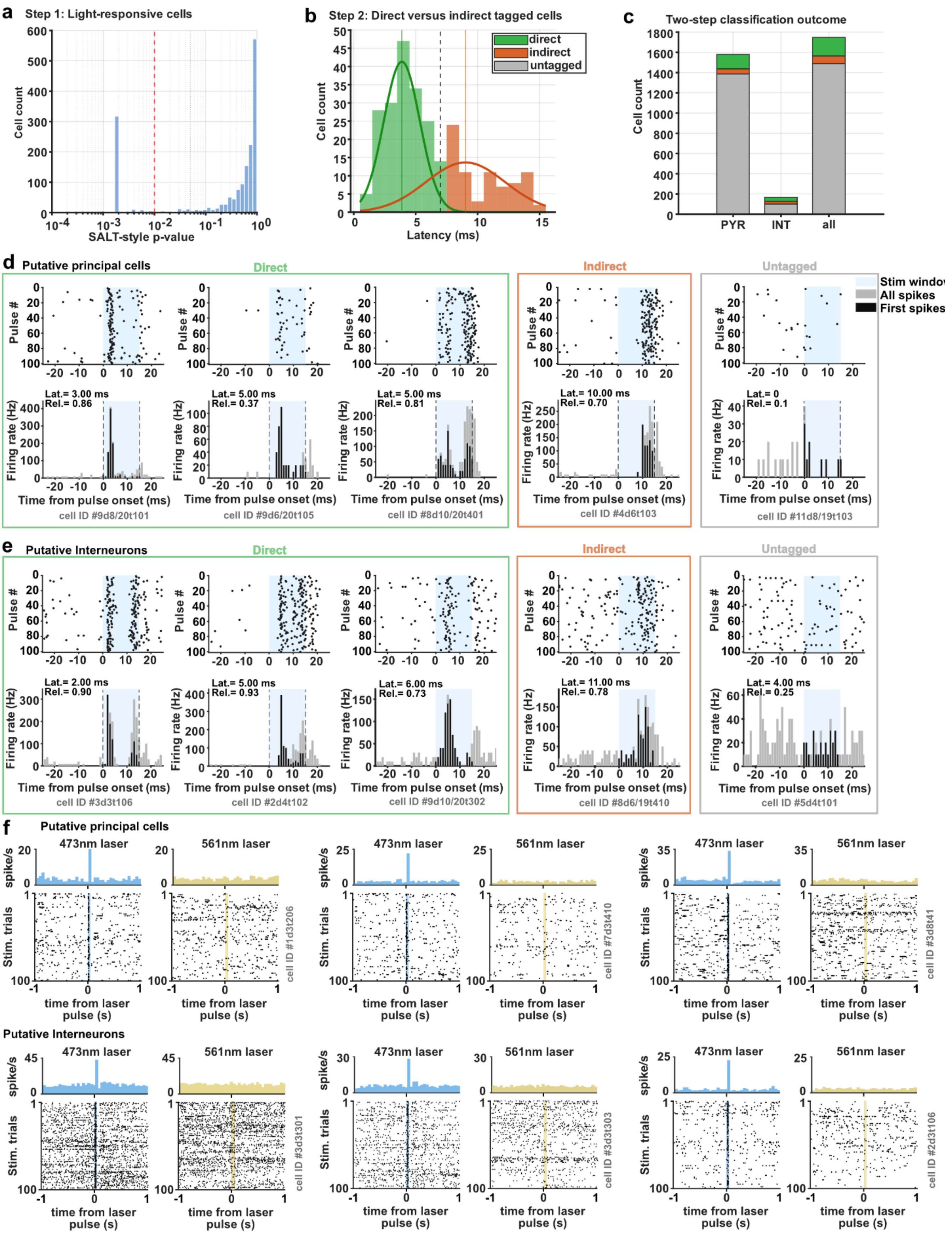


**Fig. S2. SALT based optotagging identifies directly tagged cfos+ units and separates them from indirect light responses.**

(a) Step 1 of the optotagging classification. Light responsive cells were identified using the stimulus associated spike latency test (SALT). The histogram shows SALT P values across recorded units, and the dashed red line indicates the significance threshold used to identify light responsive cells. (b) Step 2 of the classification. Light responsive units were separated into directly optotagged and indirectly activated cells based on first spike latency after light pulse onset. First spikes within the 15 ms stimulation window were used to calculate the latency, and the mode of first spike latencies across trials was used rather than the mean latency or all spikes, to avoid later secondary spikes biasing latency estimates. Cells with first spike latency above the 7 ms cutoff were classified as indirect responses and were excluded from the directly optotagged cfos+ population. (c) Outcome of the two step classification for putative principal cells, putative interneurons, and all recorded units, showing directly optotagged cells, indirect light responsive cells, and untagged cells. (d) Example putative principal cells classified as directly optotagged, indirectly light responsive, or untagged. Raster plots and peri stimulus time histograms are aligned to light pulse onset. The stimulation window is shaded, all spikes are shown in gray, and first spikes used for latency estimation are shown in black. (e) Example putative interneurons classified as directly optotagged, indirectly light responsive, or untagged, shown as in (d). (f) Control optotagging sessions using 473 nm and 561 nm laser stimulation. Directly optotagged examples showed time locked responses to 473 nm light but not to 561 nm control stimulation, supporting wavelength specific activation of ChR2 expressing cfos tagged cells.

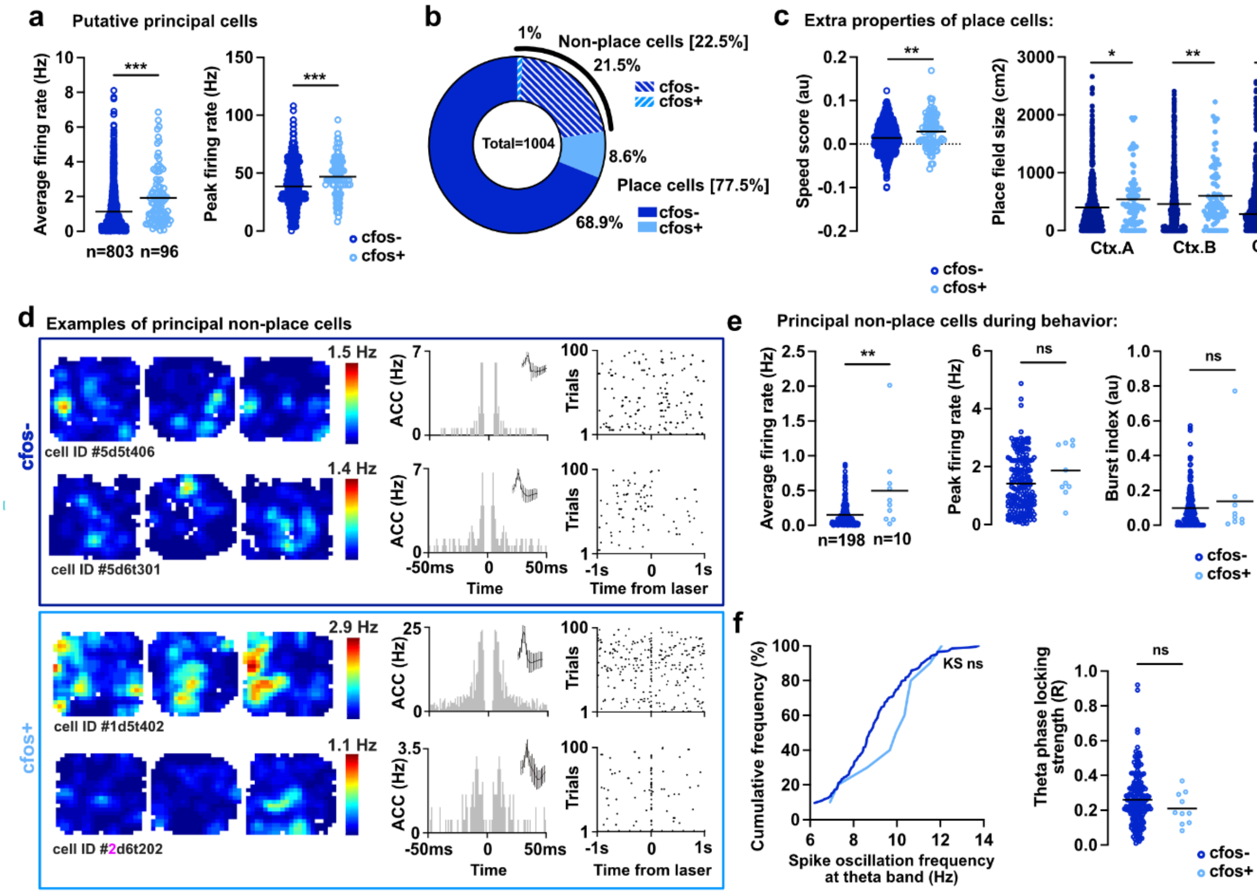


**Fig. S3. Additional characterization of cFos-tagged principal cells and principal non-place cells during sequential context exposure.**

(a) Average firing rate and temporal peak firing rate of untagged (cfos−) and cFos-tagged (cfos+) putative principal cells during behavior. (b) Distribution of putative principal cells into place-cell and non-place-cell populations, separated by cfos-tagging status. Percentages indicate the fraction of all putative principal cells in each category. (c) Additional properties of cfos− and cfos+ place cells, revealed a higher speed score and larger place-field size across Context A, Context B, and the revisit to Context A (A′). (d) Example cfos− and cfos+ principal non-place cells. For each cell, spatial firing maps are shown across Context A, Context B, and A′, together with autocorrelograms, spike waveforms, and optotagging responses. (e) Firing properties of putative principal non-place cells during behavior, including average firing rate, peak firing rate, and burst index. (f) Theta-related firing properties of putative principal non-place cells. Left, cumulative distribution of intrinsic spike oscillation frequencies in the theta band. Right, theta phase-locking strength. In a, c, e, and f, circles indicate individual cells and horizontal lines indicate the mean. ns: P > 0.05, * P < 0.05, ** P < 0.01, *** P < 0.001.

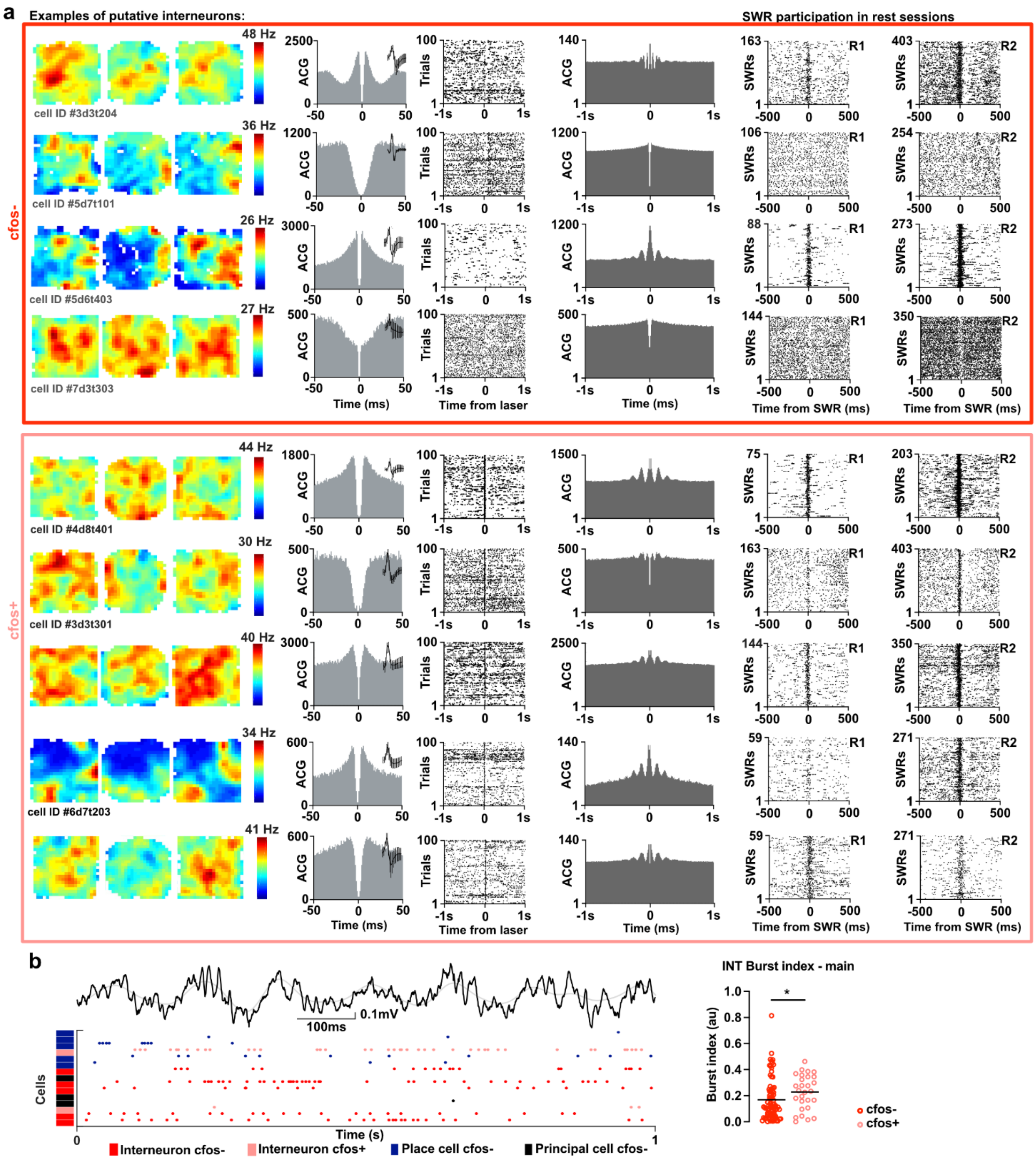


**Fig. S4. Additional examples of cFos-tagged putative interneurons and their recruitment during rest-associated SWRs.** (a) Example untagged (cfos−) and cFos-tagged (cfos+) putative interneurons recorded during sequential exposure to Context A, Context B, and the revisit to Context A (A′). For each cell, spatial firing maps across the three context exposures are shown together with autocorrelograms, spike waveforms, and optotagging responses aligned to laser stimulation. (b) Example local field potential trace with simultaneous spiking of cfos− and cfos+ putative interneurons, cfos− place cells, and cfos− principal non-place cells, together with quantification of burst index in cfos− and cfos+ putative interneurons. * P < 0.05

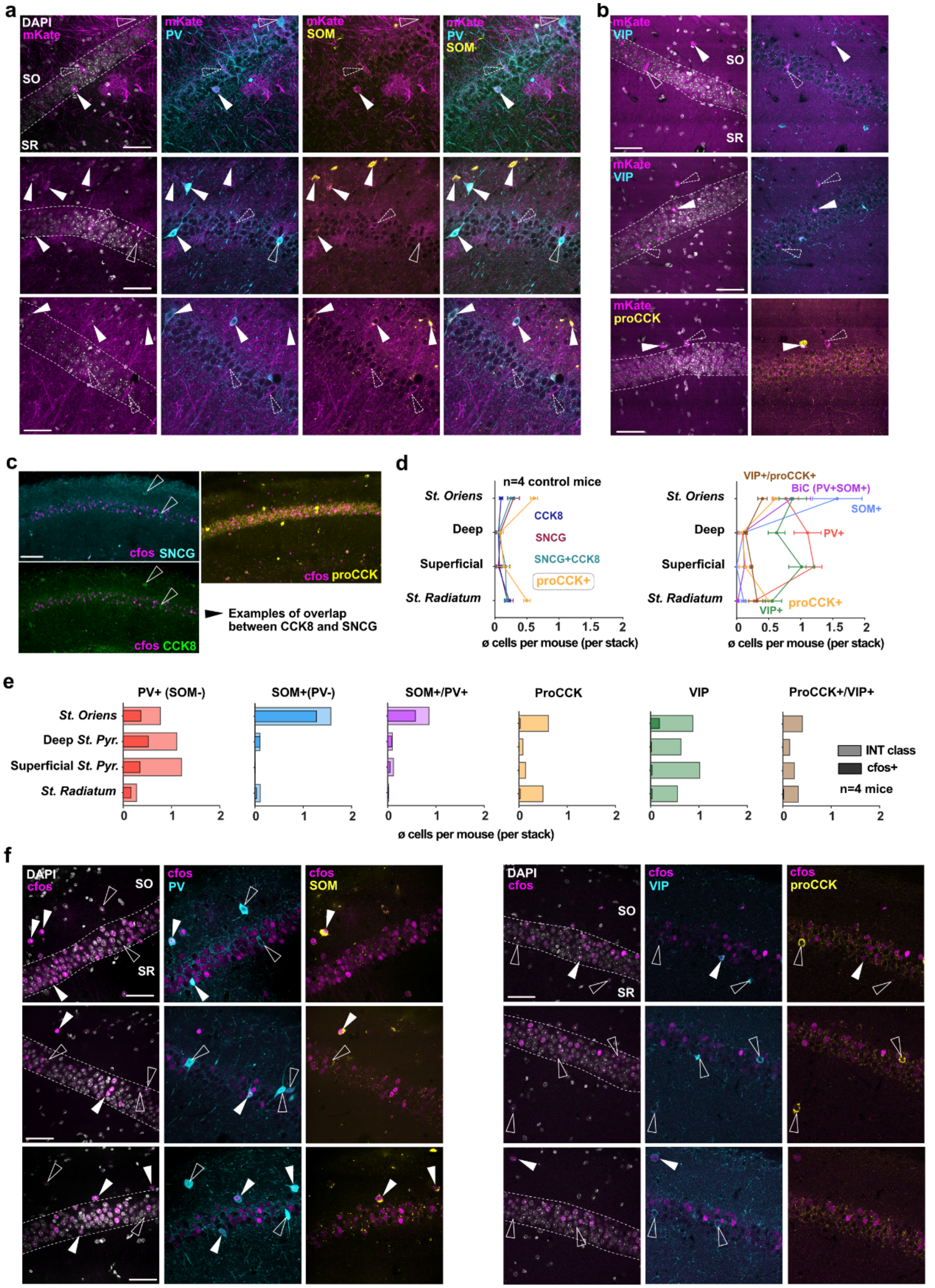


**Fig. S5. Immunohistochemical characterization of cFos-tagged and endogenous cFos-positive CA1 interneuron populations after spatial context exposure.**
(a) Representative confocal images from cFos-tTA mice after spatial context exposure, showing mKate/RFP+ cells in relation to

parvalbumin (PV) and somatostatin (SOM) immunoreactivity across CA1 layers.
(b) Representative confocal images from cFos-tTA mice showing mKate/RFP+ cells in relation to vasoactive intestinal peptide (VIP) and procholecystokinin (proCCK) immunoreactivity, illustrating mostly absence in overlap of mKate/RFP+ cells with these interneuron markers. (c) Representative confocal images from control mice stained for endogenous cFos together with SNCG, CCK8, or proCCK. Example fields illustrate the relationship between CCK-related marker expression, SNCG labeling, and endogenous cFos expression after spatial context exposure. (d) Layer-wise distribution of interneuron marker-defined cell classes across CA1. Traces show the mean number of marker-defined cells per mouse per confocal stack in stratum oriens, deep stratum pyramidale, superficial stratum pyramidale, and stratum radiatum. (e) Quantification of endogenous cFos+ cells within marker-defined interneuron classes across CA1 layers in control mice after spatial context exposure. Bars show the mean number of all marker-defined interneurons and cFos+ marker-defined interneurons per mouse per confocal stack. (f) Representative confocal examples from control mice showing endogenous cFos+ cells in relation to PV, SOM, VIP, or proCCK immunoreactivity. Filled arrowhead: cells positive for interneuron marker and mKate or cFos. Open arrowhead: cells positive for interneuron marker, but negative for mKate or cFos. Dashed arrowhead: positive for mKate or cFos, but negative for interneuron marker. Scale Bars (a,b,f): 50µm and (b): 100µm

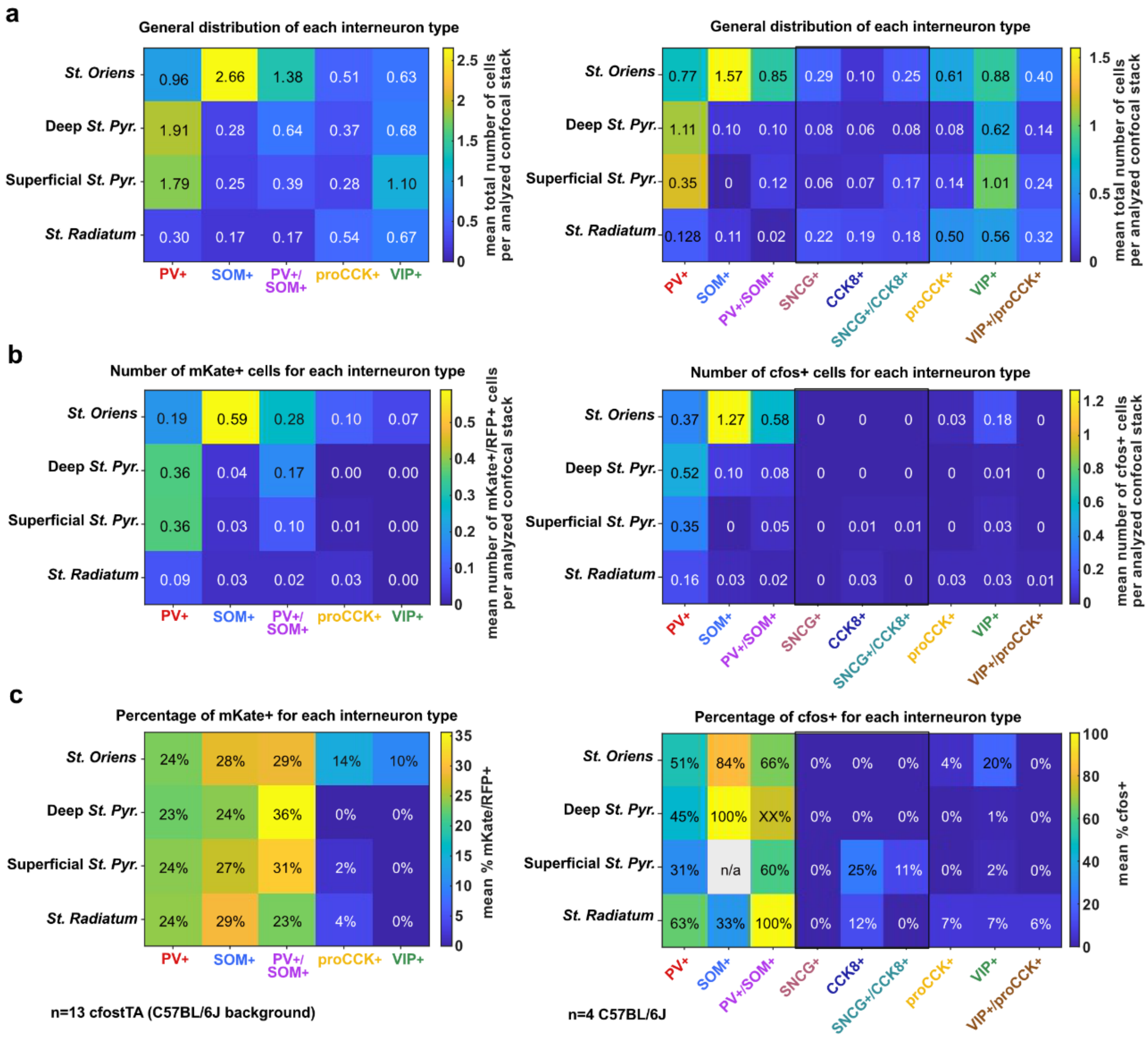


**Fig. S6. Immunohistochemical characterization of cFos-tagged CA1 cells after spatial context exposure.**
(a) Distribution of interneuron marker-defined cell classes across CA1 layers. Heat maps show the mean number of cells per analyzed slice/stack per mouse for PV+, SOM+, PV+/SOM+ double-positive, proCCK+, and VIP+ cells in stratum oriens, deep stratum pyramidale, superficial stratum pyramidale, and stratum radiatum. This analysis defines the anatomical availability of each interneuron class across CA1 layers. (b) Number of mKate/RFP+ cells within each interneuron class and layer in cFos-tTA mice after the spatial context exposure and cFos-tagging protocol. mKate/RFP+ cells were detected within PV+, SOM+, and PV+/SOM+ populations, whereas overlap with proCCK+ and VIP+ populations was low or absent. (c) Fraction of each interneuron class that was mKate/RFP+, calculated as the number of mKate/RFP+ cells within a given marker-defined class divided by the total number of cells in that class. This percentage-based analysis shows that PV+, SOM+, and PV+/SOM+ interneurons contained a relatively higher fraction of mKate/RFP+ cells than proCCK+ or VIP+ interneurons, indicating selective recruitment of these interneuron classes into the cFos-tagged CA1 population.
Right panels show the corresponding analysis for endogenous cFos expression after spatial context exposure. Similar to the mKate/RFP-based tagging analysis, endogenous cFos labeling was enriched in PV+, SOM+, and PV+/SOM+ interneuron classes, whereas proCCK+ and VIP+ cells showed little or no cFos expression. Together, these data indicate that spatial context exposure recruits a selective subset of CA1 interneurons, prominently including PV and SOM classes, into the cFos-expressing population rather than uniformly labeling all interneuron types.

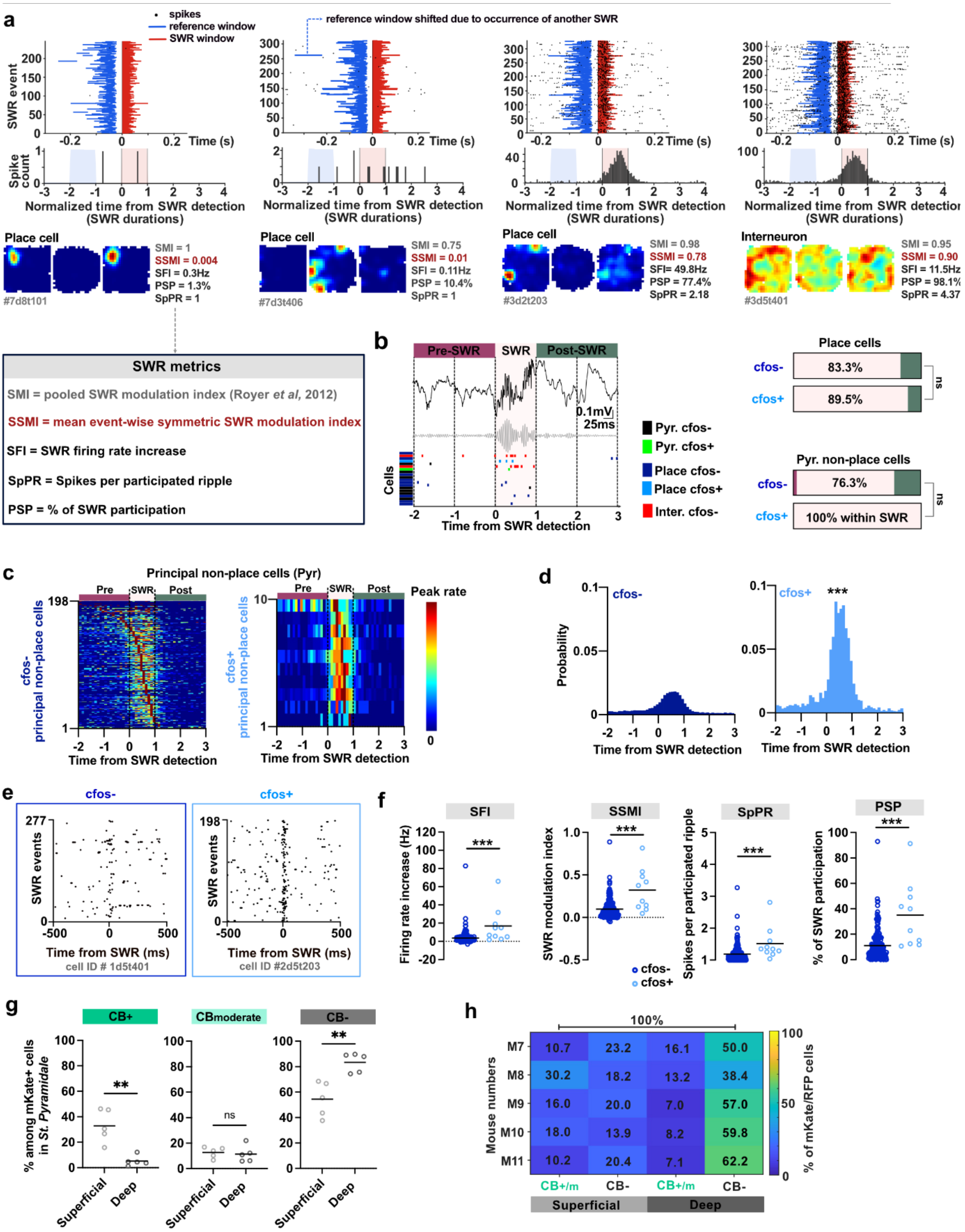


**Fig. S7. Additional examples and analyses of SWR recruitment and cFos-tagged CA1 cell identity.**
(a) Example units illustrating SWR recruitment metrics used in Fig. 4, including three place cells with corresponding open-field rate maps from OF1-OF3 and one putative interneuron. For each example cell, SWR-associated activity is summarized by the symmetric SWR modulation index (SSMI), SWR firing rate increase (SFI), percentage of SWR participation (PSP), and spikes per participated ripple (SpPR). (b) Example SWR-aligned firing profile and quantification of whether individual place and non-place principal cells reached their peak firing before, during, or after the SWR window. (c) SWR-aligned firing rate heat maps of individual untagged and cFos-tagged non-place principal cells during post-behavior rest, sorted by the relative time of their peak

firing rate, analogous to the place-cell heat maps shown in Fig. 4b. Rows indicate individual cells and columns indicate normalized time relative to SWR detection. (d) Probability distributions of SWR-associated population activity for untagged and cFos-tagged non-place principal cells, analogous to Fig. 4c. (e) Example raster plots of individual untagged and cFos-tagged non-place principal cells across detected SWR events, centered at SWR detection, analogous to the place-cell examples in Fig. 4d. (f) Quantification of SWR recruitment metrics in non-place principal cells, including SFI, SSMI, and SpPR, analogous to Fig. 4f. (g) Quantification of CB immunoreactivity among mKate/RFP+ cells in superficial and deep CA1 stratum pyramidale. Strongly CB-positive mKate/RFP+ cells were enriched in superficial CA1 and were rare in deep CA1, whereas CB-negative mKate/RFP+ cells were enriched in deep CA1. Moderately CB-positive cells were observed at similar proportions in superficial and deep CA1. (h) Complementary mouse-wise quantification of the same mKate/RFP+ cells across all radial/CB-defined groups, normalized together to 100% per mouse. In contrast to Fig. 4k, where CB+ and CB− fractions are normalized separately within superficial and deep CA1, this representation shows the overall allocation of tagged cells across superficial CB+, superficial CB−, deep CB+, and deep CB− groups. Circles indicate individual cells where applicable, and horizontal lines indicate the mean. For mouse-wise quantifications, each dot or connected line represents one mouse. Statistical annotations as in the figure.
ns: P > 0.05, ** P < 0.01, *** P < 0.001.

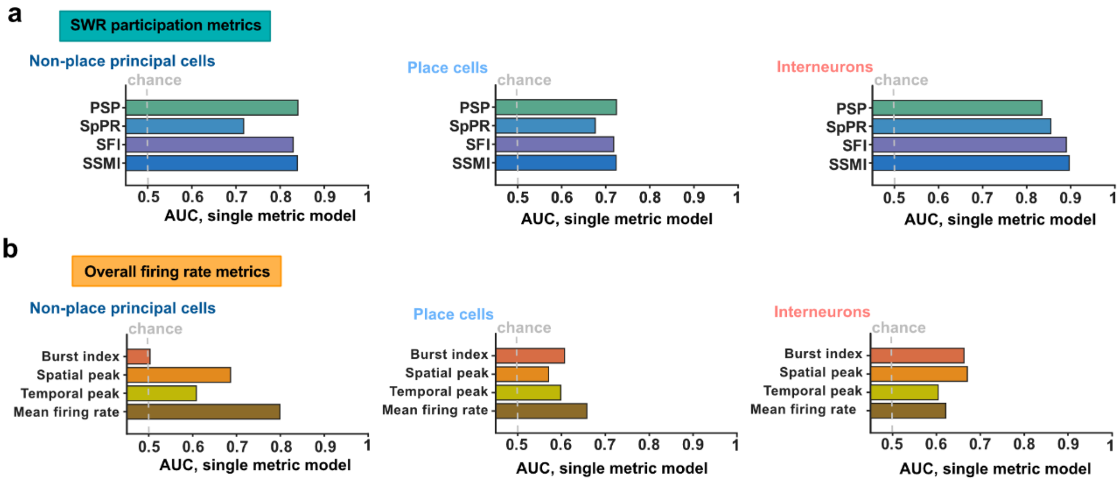


**Fig. S8. Single-metric predictive value of SWR participation and firing-rate features for cFos-tagged cell identity.**
(a) Area under the receiver operating characteristic curve (AUC) from single-metric logistic regression models using individual SWR participation metrics to predict directly optotagged cFos-tagged identity. Metrics included percentage of SWR participation (PSP), spikes per participated ripple (SpPR), SWR firing increase (SFI), and symmetric SWR modulation index (SSMI). (b) AUC from single-metric logistic regression models using individual firing-rate-related metrics, including burst index, spatial peak firing rate, temporal peak firing rate, and mean firing rate. Models were fit separately for non-place principal cells, place cells, and putative interneurons. Dashed lines indicate chance-level discrimination at AUC = 0.5.

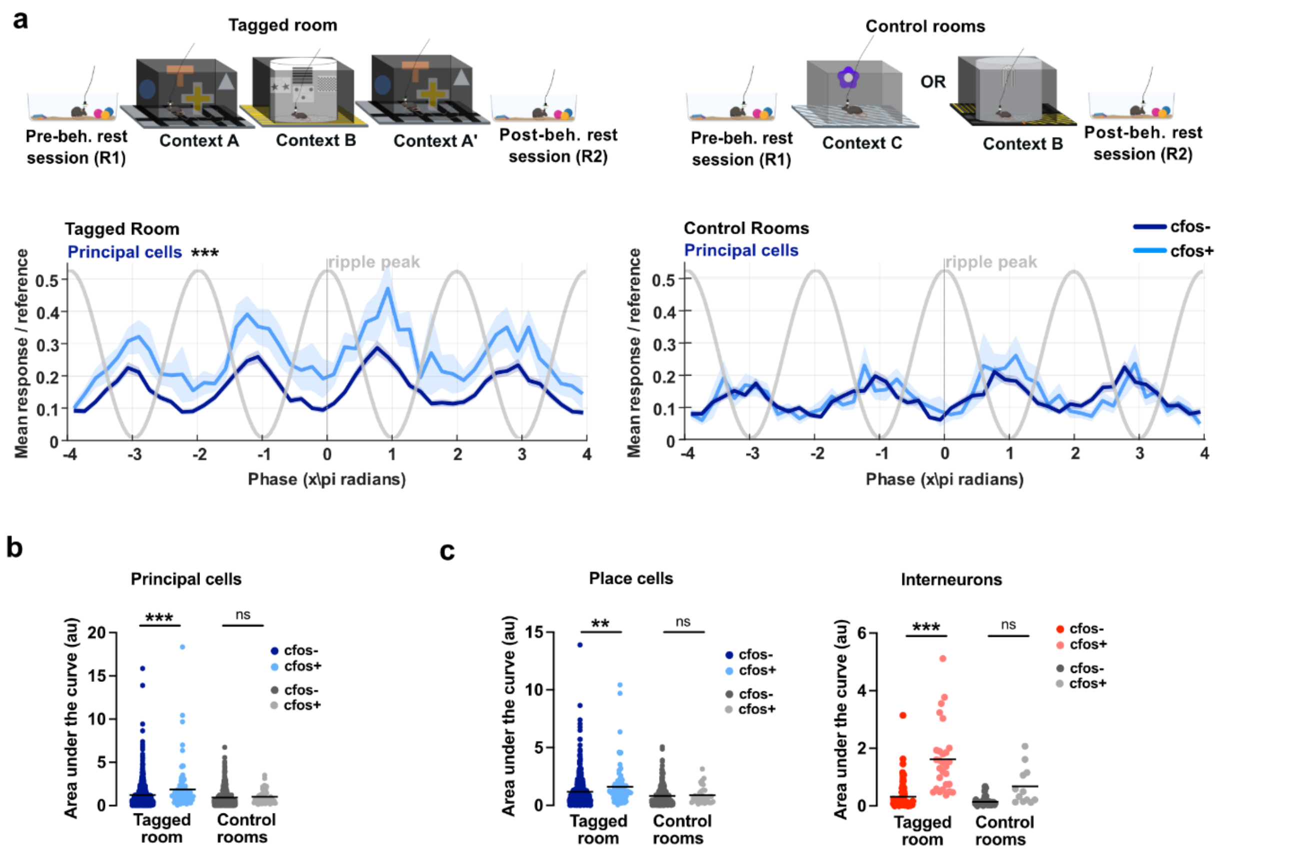


**Fig. S9. Comparison of SWR phase resolved recruitment in cFos defined cell populations between tagged and control rooms.**

(a) Schematic of the rest sessions in the tagged and control rooms, together with SWR center aligned phase response profiles for putative principal cells. Spike phases were aligned to the center ripple peak and plotted across multiple ripple cycles surrounding the SWR center. Curves show the mean reference normalized response of cFos positive and cFos negative cells; shaded regions indicate error across animals. Grey traces indicate the ripple phase.

(b) Quantification of SWR phase resolved recruitment in principal cells. Absolute AUC was calculated from each cell's reference normalized, center aligned phase response within predefined windows around the expected ripple cycle peaks, and compared between cFos-tagged and untagged cells in the tagged and control rooms.

(c) Same AUC-based quantification as in (b), shown separately for place cells and putative interneurons. cFos positive cells showed increased phase resolved recruitment in rest sessions from the tagged room, whereas this difference was reduced or absent in rest sessions from the control rooms. Dots indicate individual cells and horizontal lines indicate group summaries. Significance was assessed by ANOVA with multiple comparison testing as indicated. * $P < 0.05$, ** $P < 0.01$, *** $P < 0.001$.

**Supplementary Table 1**. Statistical analyses of cFos-tagged and untagged CA1 principal-cell composition, spatial coding, and theta-related properties. Related to Figs. 1, 2, and S3.

| Figure panel | Graph description | n cfos- | n cfos+ | Mean ± SEM cfos- | Mean ± SEM cfos+ | Stat. test | Stat. metrics | P= | Sig |
|---|---|---|---|---|---|---|---|---|---|
| Fig.1i | Putative principal cells | 803 | 96 | 90.9 % | 77.4% | Binomial test: cfos+/- vs expected (total) | | cfos-: 0.1147 | ns |
| | Putative interneurons | 80 | 28 | 9.1 % | 22.6% | | | cfos+: 0.0001 | *** |
| **PRINCIPAL CELLS** | | | | | | | | | |
| Fig.2a | Place cells | 605 | 86 | 75.3% | 89.6% | Binomial test: cfos+/- vs expected (total) | | cfos-: 0.3152 | ns |
| | Principal non-place cells | 198 | 10 | 24.7% | 10.4% | | | cfos+: 0.0022 | ** |
| Fig S3a | Average firing rate all principal cells (Hz) | 803 | 96 | 1.13 ± 0.045 | 1.92 ± 0.16 | MW | U=23562 | <0.0001 | *** |
| | Peak firing rate all principal cells (Hz) | 803 | 96 | 38.60 ± 0.67 | 46.88 ± 1.77 | MW | U=28443 | <0.0001 | *** |
| **PLACE CELLS** | | | | | | | | | |
| Fig 2b | Average firing rate (Hz) | 605 | 86 | 1.41 ± 0.054 | 2.07 ± 0.17 | MW | U=18004 | <0.0001 | *** |
| | Peak firing rate (Hz) | 605 | 86 | 45.70 ± 0.63 | 50.14 ± 1.62 | MW | U=21300 | 0.0063 | ** |
| | Burst index (au) | 595 | 86 | 0.19 ± 0.0052 | 0.22 ± 0.0126 | MW | U=20334 | 0.0020 | ** |
| Fig. 2c | Spatial peak firing rate (Hz) | 605 | 86 | 10.18 ± 0.23 | 11.45 ± 0.61 | MW | U=22249 | 0.0297 | * |
| | Place field size ($cm^2$) | 605 | 86 | 420.2 ± 16.83 | 560.7 ± 48.86 | MW | U=20172 | 0.0007 | *** |
| | Information (bits/spike) | 605 | 86 | 1.06 ± 0.033 | 0.64 ± 0.059 | MW | U=11523 | <0.0001 | *** |
| | Place field number§: #1 | 967 | 131 | 66.0% | 58.5% | Chi-square: cfos-/cfos+ vs pooled total | | cfos-: 0.4958 | ns |
| | (across Ctx A, B, A') #2 | 365 | 58 | 25.0% | 25.9% | | | | |
| | #3 | 131 | 35 | 9.0% | 15.6% | | | cfos+: 0.0101 | * |
| Fig. 2g | Place Cell Dice score | 11 mice | 11 mice | Two-way ANOVA | | Interaction | | 0.0544 | ns |
| | | | | | | cfos class | | 0.0005 | *** |
| | | | | | | open fields comp. | | 0.0010 | ** |
| | Multiple comparisons: CtxA | | | Šídák's multiple comparisons test | | | df=57 | 0.0076 | ** |
| | Ctx B | | | | | | df=57 | >0.9999 | ns |
| | CtxA' | | | | | | df=57 | 0.0184 | * |
| Fig. S3c | Speed score place cells (au) | 529 | 82 | 0.014 ± 0.0015 | 0.029 ± 0.0043 | MW | U=17215 | 0.0026 | ** |
| | Place field size (Ctx A,B, A') | 605 | 86 | Kruskal-Wallis test (Dunn's post-hoc) | | | H=27.42 | <0.0001 | *** |
| | Multiple comparisons: CtxA | | | 400.0 ± 17.5 | 539.9 ± 54.7 | Dunn | Z=2.550 | 0.0323 | * |
| | Ctx B | | | 457.5 ± 21.3 | 604.4 ± 57.9 | Dunn | Z=3.508 | 0.0014 | ** |
| | CtxA' | | | 403.1 ± 17.3 | 537.9 ± 50.7 | Dunn | Z=2.882 | 0.0143 | * |
| Fig 2i | Stability coefficient | 603 | 86 | 0.79 ± 0.093 | 0.66 ± 0.025 | MW | U=22257 | 0.0317 | * |
| | Remapping coefficient | 603 | 86 | 0.11 ± 0.0093 | 0.14 ± 0.036 | MW | U=24314 | 0.3498 | ns |
| Fig 2k | Spike oscillation frequency at the Theta band (Hz) | 605 | 86 | 8.88 ± 0.048 | 8.83 ± 0.10 | MW | U=25609 | 0.8149 | ns |
| | | | | | | KS | D=0.1154 | 0.2684 | ns |
| Fig 2l | Theta phase preference | 605 | 86 | 223.70° | 220.02° | Kuiper perm. | KV= 0.1145 | 0.7798 | ns |
| Fig 2m | Theta phase locking strength (R) | 605 | 86 | 0.1657 ± 0.0037 | 0.1565 ± 0.0072 | MW | U=25348 | 0.7004 | ns |
| **NON-PLACE CELLS** | | | | | | | | | |
| Fig S3e | Average firing rate (Hz) | 198 | 10 | 0.15 ± 0.013 | 0.50 ± 0.014 | MW | U=486 | 0.0054 | ** |
| | Peak firing rate (Hz) | 198 | 10 | 1.41 ± 0.067 | 1.87 ± 0.28 | MW | U=682 | 0.0982 | ns |
| | Burst index (au) | 198 | 10 | 0.099 ± 0.0087 | 0.14 ± 0.081 | MW | U=764 | 0.9961 | ns |
| Fig S3f | Spike oscillation frequency at the Theta band (Hz) | 198 | 10 | 8.87 ± 0.12 | 9.75 ± 0.53 | MW | U=675.5 | 0.0907 | ns |
| | | | | | | KS | D=0.4020 | 0.0922 | ns |
| | Theta phase locking strength (R) | 198 | 10 | 0.26 ± 0.011 | 0.21 ± 0.029 | MW | U=721 | 0.2750 | ns |

§ = n is actual number of place fields in context A, B and A'; **MW**= Mann-Whitney test; **KS** = Kolmogorov-Smirnov test; Dunn's = post-hoc test for multiple comparisons; **Kuiper Perm.** = two-sample Kuiper permutation test

**Supplementary Table 2.** Statistical analyses of cFos-tagged and untagged CA1 interneuron firing, speed modulation, and theta coupling. Related to Figs. 3 and S4.

| Figure panel | Graph description | n cfos- | n cfos+ | Mean ± SEM cfos- | Mean ± SEM cfos+ | Stat. test | Stat. metrics | P= | Sig |
|---|---|---|---|---|---|---|---|---|---|
| **INTERNEURONS** | | | | | | | | | |
| Fig. 3c | Average firing rate (Hz) | 80 | 28 | 16.53 ± 1.13 | 18.50 ± 1.64 | MW | U=909 | 0.1407 | ns |
| | Peak firing rate (Hz) | 80 | 28 | 94.00 ± 3.66 | 104.10 ± 6.86 | MW | U=933.5 | 0.1923 | ns |
| | Speed score (au) | 80 | 28 | 0.040 ± 0.010 | 0.14 ± 0.016 | MW | U=468 | <0.0001 | *** |
| Fig. S4b | Bust index (au) | 80 | 28 | 0.17 ± 0.018 | 0.23 ± 0.025 | MW | U=802 | 0.0254 | * |
| Fig. 3e | Spike oscillation frequency at the Theta band (Hz) | 80 | 28 | 7.68 ± 0.14 | 8.30 ± 0.18 | MW | U=865 | 0.0732 | ns |
| | | | | | | KS | D=0.3411 | 0.0160 | * |
| Fig. 3f | Theta phase preference | 80 | 28 | 180.32° | 220.95° | Kuiper perm. | KV= 0.4412 | 0.00380 | ns |
| Fig. 3g | Theta phase locking strength (R) | 80 | 28 | 0.12 ± 0.081 | 0.20 ± 0.068 | MW | U=470 | <0.0001 | * |
| **MW**= Mann-Whitney test; **KS** = Kolmogorov-Smirnov test; **Kuiper Perm.** = two-sample Kuiper permutation test | | | | | | | | | |

**Supplementary Table 3.** Statistical analyses of SWR recruitment, co-recruitment, and anatomical allocation of cFos-tagged CA1 cell populations. Related to Figs. 4, 5, and S7.

| Figure panel | Graph description | n cfos- | n cfos+ | Mean ± SEM cfos- | Mean ± SEM cfos+ | Stat. test | Stat. metrics | P= | Sig |
|---|---|---|---|---|---|---|---|---|---|
| **PLACE CELLS** | | | | | | | | | |
| Fig. S7b | SWR related spiking - before<br>during<br>after | 605 | 86 | 16.0%<br>82.3%<br>0.7% | 10.5%<br>89.5%<br>0% | Chi-square | $X^2$=2.741 df=2 | 0.2540 | ns |
| Fig. 4c | Probability of spiking | 605 | 86 | n/a | | KS | D=0.0423 | <0.0001 | *** |
| Fig. 4f | SWR metrics SFI | 605 | 86 | 7.06 ± 0.40 | 13.88 ± 1.43 | MW | U=15566 | <0.0001 | *** |
| | SSMI | 605 | 86 | 0.17 ± 0.0060 | 0.29 ± 0.021 | MW | U=15194 | <0.0001 | *** |
| | SpPR | 605 | 86 | 1.48 ± 0.035 | 1.27 ± 0.012 | MW | U=17424 | <0.0001 | *** |
| Fig. 4g | PSP | 605 | 86 | 20.46 ± 0.68 | 34.47 ± 2.30 | MW | U=15109 | <0.0001 | *** |
| **NON-PLACE PRINCIPAL CELLS** | | | | | | | | | |
| Fig. S7b | SWR related spiking - before<br>during<br>after | 198 | 10 | 20.7%<br>76.3%<br>3.0% | 0%<br>100%<br>0% | Chi-square | $X^2$=2.330 df=2 | 0.3119 | ns |
| Fig. S7d | Probability of spiking | 198 | 10 | n/a | | KS | D=0.0767 | <0.0001 | *** |
| Fig. S7f | SWR metrics SFI | 198 | 10 | 3.70 ± 0.48 | 16.77 ± 6.18 | MW | U=325 | 0.0001 | *** |
| | SSMI | 198 | 10 | 0.097 ± 0.0077 | 0.32 ± 0.078 | MW | U=339 | 0.0002 | *** |
| | SpPR | 198 | 10 | 1.19 ± 0.019 | 1.52 ± 0.16 | MW | U=381.5 | 0.0008 | *** |
| | PSP | 198 | 10 | 11.07 ± 0.81 | 35.07 ± 8.21 | MW | U=293 | <0.0001 | *** |
| Fig. 4h | % mKate cells in CA1 St. Pyramidale | 13 mice | | Deep: 61.60 ± 1.81 | Superficial: 38.40 ± 1.81 | MW | 0 | <0.0001 | *** |
| Fig. 4j | % overlap of mKate cells in CA1 St. Pyramidale with Calbindin signal (CB) | 5 mice | | CB-: 72.65 ± 4.34 | CB+/mod.: 27.35 ± 4.35 | MW | 0 | 0.0079 | ** |
| Fig. S7g | Distribution of mKate+ cells in St. Pyramidale | | | Superficial | Deep | | | | |
| | CB positive | 5mice | | 32.92 ± 5.79 | 5.22 ± 1.84 | MW | 0 | 0.0079 | ** |
| | CB moderate | | | 12.70 ± 1.88 | 11.42 ± 2.91 | MW | 0 | 0.5476 | ns |
| | CB negative | | | 54.38 ± 6.11 | 83.36 ± 3.42 | MW | 0 | 0.0079 | ** |
| **INTERNEURONS** | | | | | | | | | |
| Fig. 5c | SWR related spiking - before<br>during<br>after | 80 | 28 | 37.5%<br>42%<br>20% | 3.6%<br>96.6%<br>0% | Chi-square | $X^2$=27.15 df=2 | <0.0001 | *** |
| Fig. 5d | Probability of spiking | 80 | 28 | n/a | | KS | D=0.1090 | <0.0001 | *** |
| Fig. 5f | SWR metrics SFI | 80 | 28 | 13.83 ± 4.40 | 76.26±11.11 | MW | U=267 | <0.0001 | *** |
| | SSMI | 80 | 28 | 0.22 ± 0.037 | 0.64 ± 0.045 | MW | U=246 | <0.0001 | *** |
| | SpPR | 80 | 28 | 1.73 ± 0.15 | 3.57 ± 0.42 | MW | U=350.5 | <0.0001 | *** |
| Fig. 5g | PSP | 80 | 28 | 39.59 ± 3.19 | 78.49 ± 4.97 | MW | U=393 | <0.0001 | *** |
| Fig. 5h | Co-activity metric | n=7 mice | | cfos+>cfos- 0.44 ± 0.13 | cfos+>cfos+ 1.13 ± 0.20 | WCx | n/a | 0.0078 | ** |
| **MW**= Mann-Whitney test; **KS** = Kolmogorov-Smirnov test; **WCx.** = Wilcoxon signed-rank test | | | | | | | | | |

**Supplementary Table 4.** Statistical summary of logistic-regression models relating firing-rate and SWR-recruitment features to cFos-tagged cell identity. Related to Figs. 6 and S8.

| Figure panel | Graph description | Metric | Statistical tests | Stat. metrics | P= | Sig |
|---|---|---|---|---|---|---|
| Fig. 6b | AUC permutation test | Observed AUC | Null AUC mean [95% null Conf. interval] | Permutations | | |
| | Non-place principal cells | 0.842 | 0.626 [0.495-0.762] | 10000 | 0.0008 | *** |
| | Place cells | 0.723 | 0.539 [0.498- 0.589] | 10000 | 0.0001 | *** |
| | Interneurons | 0.883 | 0.577 [0.500 – 0.673] | 10000 | 0.0001 | *** |
| Fig. 6d | | Odds ratio | Confidence interval | | | |
| | NON-PLACE PRINCIPAL CELLS | | | | | |
| | Firing Rate metrics | 1.10 | 0.50 – 2.39 | | 0.818 | ns |
| | SWR metrics | 3.20 | 1.60 – 6.41 | | <0.0001 | *** |
| | PLACE CELLS | | | | | |
| | Firing Rate metrics | 0.99 | 0.78 – 1.27 | | 0.964 | ns |
| | SWR metrics | 1.97 | 1.58 – 2.46 | | <0.0001 | *** |
| | INTERNEURONS | | | | | |
| | Firing Rate metrics | 0.67 | 0.34 – 1.35 | | 0.257 | ns |
| | SWR metrics | 5.23 | 2.47 – 11.17 | | <0.0001 | *** |
| Fig. 6e | | Model fit lost | Chi-square | | | |
| | NON-PLACE PRINCIPAL CELLS | | | | | |
| | Dropped block: Firing Rate metrics | 0.44% | 0.085 | | 0.771 | ns |
| | Dropped block: SWR metrics | 65.96% | 12.676 | | 0.0004 | *** |
| | PLACE CELLS | | | | | |
| | Dropped block: Firing Rate metrics | 0.39% | 0.192 | | 0.662 | ns |
| | Dropped block: SWR metrics | 73.96% | 36.721 | | <0.0001 | *** |
| | INTERNEURONS | | | | | |
| | Dropped block: Firing Rate metrics | 4.65% | 1.612 | | 0.204 | ns |
| | Dropped block: SWR metrics | 86.87% | 30.142 | | <0.0001 | *** |
| Fig. 6f | | Odds ratio | Confidence interval | | | |
| | NON-PLACE PRINCIPAL CELLS | | | | | |
| | PSP | 3.18 | 1.79 – 5.63 | | <0.0001 | *** |
| | SpPR | 1.87 | 1.19 – 2.92 | | 0.0058 | ** |
| | SFI | 3.25 | 1.70 – 6.20 | | 0.0033 | *** |
| | SSMI | 3.19 | 1.81 – 5.63 | | <0.0001 | *** |
| | PLACE CELLS | | | | | |
| | PSP | 2.14 | 1.74 – 2.63 | | <0.0001 | *** |
| | SpPR | 1.69 | 1.39 – 2.06 | | <0.0001 | *** |
| | SFI | 1.95 | 1.60 – 2.38 | | <0.0001 | *** |
| | SSMI | 2.17 | 1.76 – 2.67 | | <0.0001 | *** |
| | INTERNEURONS | | | | | |
| | PSP | 4.67 | 2.45 – 8.90 | | 0.0009 | *** |
| | SpPR | 3.31 | 1.62 – 6.75 | | <0.0001 | *** |
| | SFI | 4.08 | 2.05 – 8.13 | | <0.0001 | *** |
| | SSMI | 7.62 | 3.38 – 17.20 | | <0.0001 | *** |

AUC = area under the curve;

**Supplementary Table 5.** Statistical analyses of context-dependent and ripple phase-resolved SWR recruitment in tagged and control rooms. Related to Figs. 7 and S9.

| Figure panel | Graph description | n cfos- | n cfos+ | Mean ± SEM cfos- | Mean ± SEM cfos+ | Stat. test | Stat. metrics | P= | Sig |
|---|---|---|---|---|---|---|---|---|---|
| **PRINCIPAL CELLS** | | | | | | | | | |
| Fig. S9b | SWR phase analysis (area under the curve) | | | Kruskal-Wallis test (Dunn's post-hoc) | | | H=48.64 | <0.0001 | |
| Fig. S9a | Multiple comparisons: Tagged room | 800 | 96 | 1.21 ± 0.048 | 1.86 ± 0.25 | Dunn | Z=3.771 | 0.0007 | *** |
| Fig. S9a | Control rooms | 623 | 47 | 0.92 ± 0.037 | 1.01 ± 0.12 | Dunn | Z=1.229 | 0.8764 | ns |
| **PLACE CELLS** | | | | | | | | | |
| Fig. S9c | SWR phase analysis (area under the curve) | | | Kruskal-Wallis test (Dunn's post-hoc) | | | H=53.10 | <0.0001 | *** |
| Fig 7e | Multiple comparisons: Tagged room | 605 | 86 | 1.12 ± 0.049 | 1.61 ± 0.18 | Dunn's | Z=3.144 | 0.0067 | ** |
| Fig 7f | Control rooms | 370 | 36 | 0.81 ± 0.041 | 0.87 ± 0.12 | Dunn's | Z=0.7375 | >0.9999 | ns |
| **INTERNEURONS** | | | | | | | | | |
| Fig. S9c | SWR phase analysis (area under the curve) | | | Kruskal-Wallis test (Dunn's post-hoc) | | | H=64.69 | <0.0001 | *** |
| Fig 7e | Multiple comparisons: Tagged room | 80 | 28 | 0.32 ± 0.054 | 1.62 ± 0.22 | Dunn's | Z=6.732 | <0.0001 | *** |
| Fig 7f | Control rooms | 48 | 12 | 0.34 ± 0.021 | 0.68 ± 0.19 | Dunn's | Z=1.718 | 0.05969 | ns |
